# Global Screening of Sentrin-Specific Protease Family Substrates in SUMOylation

**DOI:** 10.1101/2020.02.25.964072

**Authors:** Yunzhi Wang, Xiaohui Wu, Rui Ge, Lei Song, Kai Li, Sha Tian, Lili Cai, Mingwei Liu, Wenhao Shi, Guoying Yu, Bei Zhen, Yi Wang, Fuchu He, Jun Qin, Chen Ding

**Author notes:** Co-first author (Yunzhi Wang, Xiaohui Wu, Rui Ge). Correspondence (C.D.); (J.Q.); (F.H.).

## Abstract

Post-translational modification of proteins by the addition of small ubiquitin-related modifier (SUMO) is a dynamic process, in which deSUMOylation is carried out by members of the Sentrin/SUMO-specific protease (SENP) family. While identification of SUMOylation sites at global scale has made great progress, much less effort has been made on the SENP family-dependent deSUMOylation. Here we report a dataset of 3,763 high confident SUMO1 modification sites and their dependence on the 6 members of SENP proteins. Interrogation of the dataset led to the discovery that SENP3 regulates the innate immune response via deSUMOylation of DHX15 and PCBP2 and recruitment of inflammatory molecules. Collectively, this dataset presents a site-resolved network of the SUMO-SENP system, providing information for potential substrates of the SENP proteins.

## Introduction

SUMOs are small ubiquitin-like post-translational modifiers that are covalently attached to the side chain of lysine residues on the target proteins. SUMOylation plays regulatory roles in many cellular processes, including gene expression, DNA damage response, intracellular trafficking and cell cycle^1–4^. The conjugation of SUMO to proteins is accomplished by an enzymatic cascade, which includes the E1-activating enzyme SAE1/2, the E2 conjugation enzyme UBC9 and one of several E3 ligases, resulting in the formation of an isopeptide bond between the C-terminal carboxyl group of Gly in SUMO and the ε-amino group of Lys residue in the substrate protein^5^. Among the three SUMOs in human, SUMO1 shares ∼ 48% identity with SUMO2/3, whereas SUMO2 and SUMO3 are nearly identical with only three amino acid difference. Both SUMO1 and SUMO2/3 can be conjugated to a distinct and overlapping set of substrates^3,6^. SUMO2 is the predominantly expressed isoform and is indispensable for mouse embryonic development^7^. Besides, SUMOs can also form poly-SUMOylation linkages analogous to that of ubiquitin^6,8^.

SUMO conjugation is a reversible process. The removal of SUMO, also known as deSUMOylation, is catalyzed by Sentrin/SUMO-specific proteases (SENPs) that de-conjugate SUMO from the substrate proteins. Human expresses six SENPs: SENP1, 2, 3 and SENP5, 6,7. The six SENPs have been shown to reside in different subcellular localizations and process distinct substrate specificity^6,8^. SENP1 and SENP2 primarily localize in the nucleus; SENP1 is crucial for deSUMOylating SUMO1-modified proteins while SENP2 is most efficient for SUMO2/3 de-conjugation. SENP1 deSUMOylates specific target proteins to control their transcriptional activity^9–11^. SENP2 is involved in the gene expression regulation in developmental processes, where its deficiency causes cardiac defects and neurodegeneration^12,13^. SENP3 and SENP5 are enriched in the nucleolus. SENP3 functions in the ribosome biogenesis and regulates the maturation of the 28S rRNA^14^. SENP5 participates in mitochondrial fragmentation during mitosis^15^. SENP6 is the significant chain-editing enzyme and mediates multiple signaling pathways controlled by poly-SUMOylation^16^. SENP7 acts in chromatin remodeling and chromatin dynamics^16,17^. However, the global view of the substrates of different SENPs remains largely unknown.

Recently progress in mass spectrometry-based proteomics allows global discovery of post-translational modifications (PTMs), including phosphorylation, acetylation, ubiquitination and methylation^18–21^. The utilization of a His_6_-SUMO2^T90K^ mutant coupled with a diGly-Lys specific antibody greatly facilitated the identification of SUMOylated peptides. As a result, 1,002 SUMO2-modified sites were identified under normal and heat-shocked conditions^5^. Similarly, the His_10_-Lysine-deficient (K0) SUMO2 method allowed the identification of 40,765 SUMO2 sites in different stimulation conditions^22^. In contrast, only 295 SUMO1 sites were reported for the SUMO1 modification proteome^23^. We also generated a pan-SUMO1 antibody specific to SUMO1 and identified the first endogenous SUMO1 modified sites dataset with 53 high-confidence sites from mouse testis^24^. A global view of deSUMOylation by different SENPs is still not available, hindering our understanding of dynamic regulation of SUMOylation as well as the specific functions of SENPs.

Here, we employed the His_10_-SUMO1^T95R^ coupled with a diGly-Lys specific antibody strategy^23^ to map the site-specific SUMO1 modification reference proteome and identified 3,763 SUMO1 modified sites. We also measured SUMO1 modification sites that are dependent on the individual SENP members, finding that SUMO1 modified proteins that are deSUMOylated dependent on different SENPs are involved in distinct biological processes in a SENP-dependent manner, suggesting functional specificity for the SENP members. SENP3 was identified a functional role in the immune response. This study provides a rich resource for the site-specific profile of the SUMOylation/deSUMOylation in the SENP-SUMO1 systems, connecting SENPs to different cellular pathways.

## Results

### Profiling of SUMO1 modified proteome

Since it remains very challenging to directly purify and identify SUMO1-modified proteins, we employed a previously published His_10_-tagged SUMO1 mutant in which internal T (Threonine) were replaced by K (Lysine)^23,25^. Proteins that are covalently attached to His_10_-SUMO1^T95K^ mutants via their SUMOylated Lysines can be purified by Ni-NTA via His_10_ tag. Enriched His_10_-SUMO1^T95K^ mutant modified substrates then digested with Lys-C, which create diGly remnants on modified Lysines, which can be further purification by K-ε-GG antibody (**Supplementary Figure 1a**). By mutating the residue immediately preceding the diGly motif to lysine rather than arginine, and by using endoproteinase Lys-C rather than trypsin, this approach eliminated potential misidentification of sites modified by other members of the Ubl family that also contain Arginine in this position (ubiquitin, NEDD8 and ISG15).

However, we found the commercially K-ε-GG antibody can hardly enrich any diGly modified peptides generated by this method (for thirty million mutant cells less than twenty sites can be identified per biological replicate) (**Supplementary Figure 1a, Supplementary Table 1**). To maximize the number of detected SUMO1 modified sites we then changed our strategy by employing a previously reported approach to construct a His_10_-SUMO1^T95R^ mutant stable cell line^26^. The SUMO1^T95R^ mutation allows the release of the diGly remnant linked to the lysine residue in the substrate after trypsin cleavage and could also be further enriched by K-ε-GG antibody and identified with LC-MS/MS (**Supplementary Figure 1a**). And we found by using the same amount of SUMO1^T95R^ cell line, the large proportion of peptides encompassing a SUMO1 remnant-modified lysine present in the enriched samples (**Supplementary Figure 1b**). We carefully compared the number of SUMO1 modified peptides enriched by K-ε-GG antibody in two types of mutant SUMO1 cell lines, and found before K-ε-GG antibody enrichment the number of SUMO1 modified peptides detected in the two types of mutated SUMO1 cell lines were comparable (25 versus 24), yet, after K-ε-GG antibody enrichment, the number of SUMO1 modified peptides detected in the SUMO1^T95R^ mutant cell line were much higher than SUMO1 modified peptides detected in the SUMO1^T95K^ mutated cell line (598 SUMO1 modified sites per biological replicate versus 15 SUMO1 modified sites per biological replicate) (**Supplementary Figure 1c**). The abundance of SUMO1 modified peptides show the same tendency, which were much higher in SUMO1^T95R^ mutated cells than in SUMO1^T95K^ mutated cells after K-ε-GG antibody enrichment (**Supplementary Figure 1d**).

To exclude the possibility that the significant difference between the number of enriched SUMO1 modified peptides by K-ε-GG antibody was due to the two types of mutant SUMO1s’ expression level, we compared their expression pattern and level of conjugation to target proteins between two types of SUMO1 mutants, and immunoblot confirmed that the expression level of His_10_-SUMO1^T95K^ were comparable to His_10_-SUMO1^T95R^ and the level of proteins conjugated by His_10_-SUMO1^T95K^ is at the same level with proteins conjugated by His_10_-SUMO1^T95R^ (**Figure 1d**). We then compared the total abundance and number of proteins identified in two different cell lines before and after K-ε-GG antibody enrichment. As shown in supplementary figure 1e and f, even though we detected less diGly modified peptides after K-ε-GG antibody enrichment by using His_10_-SUMO1^T95K^ cell line, the total abundance of all enriched peptides (modified and unmodified), the number of total enriched proteins (modified and unmodified) were approximately the same between experiments using two types of SUMO1 mutant cell lines (**Supplementary Figure 1e, f**), which indicating the low proportion of SUMO1 modified peptides after K-ε-GG antibody enrichment is not due to the different amount of protein used for enrichment. Extracting ion chromatogram of diGly modified peptides also confirmed the higher enrichment efficiency of K-ε-GG antibody by using His_10_-SUMO1^T95R^ mutant cell lines (**Supplementary Figure 1g, h**)

**Figure 1.**
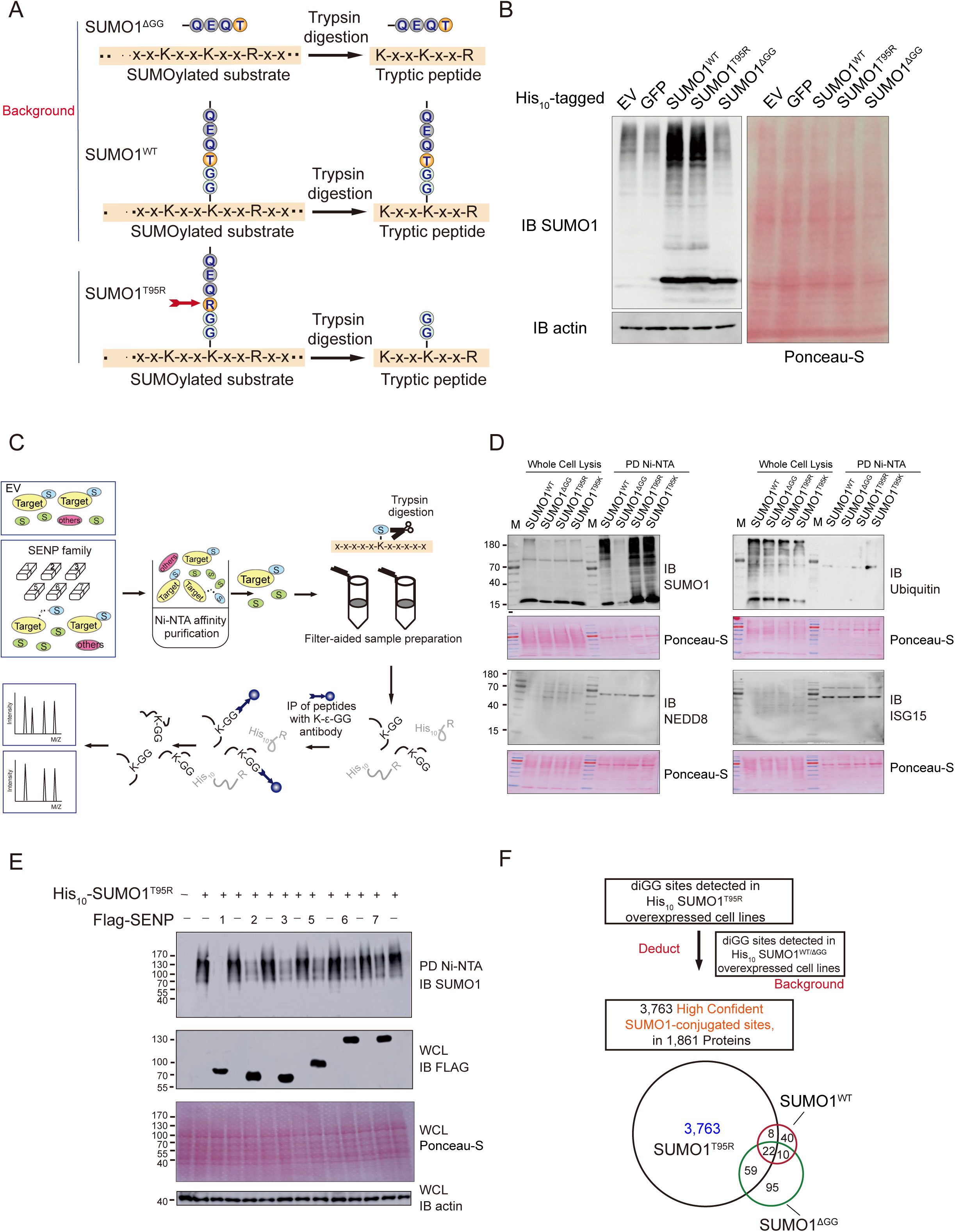
Strategy of SUMO1 modified sites immunoaffinity purification. (A) Representation of the C-terminal sequence comparison of SUMO1^WT^, SUMO1^ΔGG^ and SUMO1^T95R^ after trypsin digestion. (B) Immunoblot analysis confirmed the expressions of SUMO1-conjugated proteins in SUMO1^WT^ and SUMO1^T95R^. HEK293T cells were stably transfected with indicated plasmids and SUMO1^ΔGG^ was chosen as the negative control. Ponceau-S staining was shown as loading control. (C) Schematic overview of His_10_-SUMO1^T95R^-modified peptides identification. Stably expressing cell samples transfected with indicated plasmids were subjected to the first concentration of SUMOylated proteins and trypsin digestion, followed by peptide IP to enrich SUMO1-conjugated peptides. diGly featured peptides were subsequently exposed for LC-MS/MS identification. (D) Validation of expression patterns and level of conjugation by immunoblot analysis. His_10_-SUMO1^WT^, His_10_-SUMO1^ΔGG^, SUMO1^T95R^ and SUMO1^T95K^ stable cells were lysed and subjected to Ni-NTA pulldown assay. The presence of ubiquitin and ubiquitin-like modifier patterns were confirmed with indicated antibodies. (E) Detection of SUMOylation patterns after transfection of SENP family members. SUMOylated proteins from His_10_-SUMO1^T95R^ stably expressed cells with EV and Flag-tagged SENP family members transfected were pulled down and analyzed by immunoblot using anti-SUMO1, anti-Flag or anti-Actin antibodies. Ponceau-S staining was shown as loading control. (F) Schematic workflow of the deduction strategy for constructing SUMO1-modified proteome. Venn plot indicated the number of overlapped diGly sites detected in His_10_-SUMO1^ΔGG^, SUMO1^T95R^ and SUMO1^T95K^ stable cells.

To overcome the limitation of SUMO1^T95K^ mutation, and to construct the SUMO1 modified proteome in a large-scale, we chose SUMO1^T95R^ mutation cooperated with Tryptic digestion and K-ε-GG specific antibody enrichment strategy^23,26^ to map the site-specific SUMO1 modification reference proteome (**Figure 1c**). The His_10_-SUMO1^T95R^ expressing cells exhibited comparable SUMOylation patterns with the wild type SUMO1 (**Figure 1b**).

The approach using His_10_-tagged SUMO1^T95R^ mutant cell line has a limitation that we have to rule out the contamination by ubiquitin and ubiquitin-like modifiers such as ISG15, NEDD8, which also generated diGly remnant after tryptic digestion that can be recognized by K-ε-GG antibody. Under this consideration, we firstly performed Western Blotting analysis on SUMO1, ubiquitin, ISG15, and NEDD8, in His_10_-SUMO1^T95R^ to detect their expression pattern and level of their conjugation to target protein. The results show SUMO1^T95R^ has high conjugation level to target protein after Ni-NTA pull down, and other ubl modifiers were almost undetectable, which conform the specificity of Ni-NTA pull down method in enriching SUMOylated proteins (**Figure 1d**). Even though the immunoblotting analysis proves Ni-NTA pull down strategy is specific enough to rule out the ubiquitination and other ubl modifiers, we set His_10_-SUMO1^ΔGG^ mutant that could not be conjugated to the substrates and His_10_-SUMO1^WT^ which could not cleave by trypsin to generate diGly remnant as control, and exclude the diGly modified sites detected in those cells to build high confidence SUMO1 modification dataset (**Figure 1a, f**).

Importantly, since SUMO1, ubiquitination, ubls are all linked to the ε-amino group of lysine, the strict deduction strategy, we applied, may lead to some potential SUMO1 modification sites to be excluded. By comparing among the overlapped diGly modified peptides detected in His_10_-SUMO1^T95R^ cells, in His_10_-SUMO1^ΔGG^ cells and in His_10_-SUMO1^WT^ cells, we found that the abundance of those overlapped diGly modified peptides detected in His_10_-SUMO1^T95R^ cells were ten times higher than the abundance of those diGly modified peptides detected in His_10_-SUMO1^ΔGG^ cells or in His_10_-SUMO1^WT^ cells. This finding shows the abundance of those overlapped diGly modified peptides detected in His_10_-SUMO1^T95R^ cells were contributed mainly by SUMO1^T95R^ modified peptide. The observation was also confirmed by manually extracting XIC (extracted ion chromatogram) **(Supplementary Figure 2a)**. Therefore, our deduction strategy is strict enough to ensure the specificity of our SUMO1 dataset (**Supplementary Figure 2b**).

To establish the SUMOylation/deSUMOylation network of the SUMO1-SENP systems, we then individually over-expressed six SENPs in the His_10_-SUMO1^T95R^ stable cell line and used the empty vehicle (EV) transfected cells as the control for the SENP groups. As shown in figure 1e, while a strong SUMOylation signal was observed in the His_10_-SUMO1^T95R^ stable cell line, expressing SENP family members led to the reduction of SUMOylation proteome to various extents, with SENP1 giving rise to the most reduction, and SENP5-7 the least reduction (**Figure 1e**).

To maximize the SUMO1 modification proteome that can be reproducibly identified with our approach, we carried out 10 biological repeats for the SUMO1^T95R^-conjugated proteome. Our data revealed that the number of SUMO1^T95R^ modified peptides approached a plateau after 10 biological repeats (**Figure 2a**). We characterized 2,198 SUMO1 modified sites that represent 1,066 SUMOylated proteins (**Supplementary Table 1**). To identify SENP-dependent SUMO1 modification, we carried out 6 biological repeats for each SENP member. While a decrease in SUMO1 modification sites is expected when SENPs were expressed, new SUMO1 modification sites were also identified. Overexpression of SENPs in the SUMO1^T95R^-SENP systems led to the identification of 1,643 additional SUMO1 sites on 818 SUMOylated proteins, ranging from 119 SUMOylated sites of SENP2 to 209 SUMOylated sites of SENP6-overexpressing cells (**Supplementary Figure 3b**). Our data suggest that overexpression of SENPs can induce alterations in SUMO1 modification levels. Therefore, we defined the SUMO1^T95R^ modified sites as the reference SUMO1 map, and the SUMO1^T95R^-SENPs sites as the dynamic SUMO1 map.

**Figure 2.**
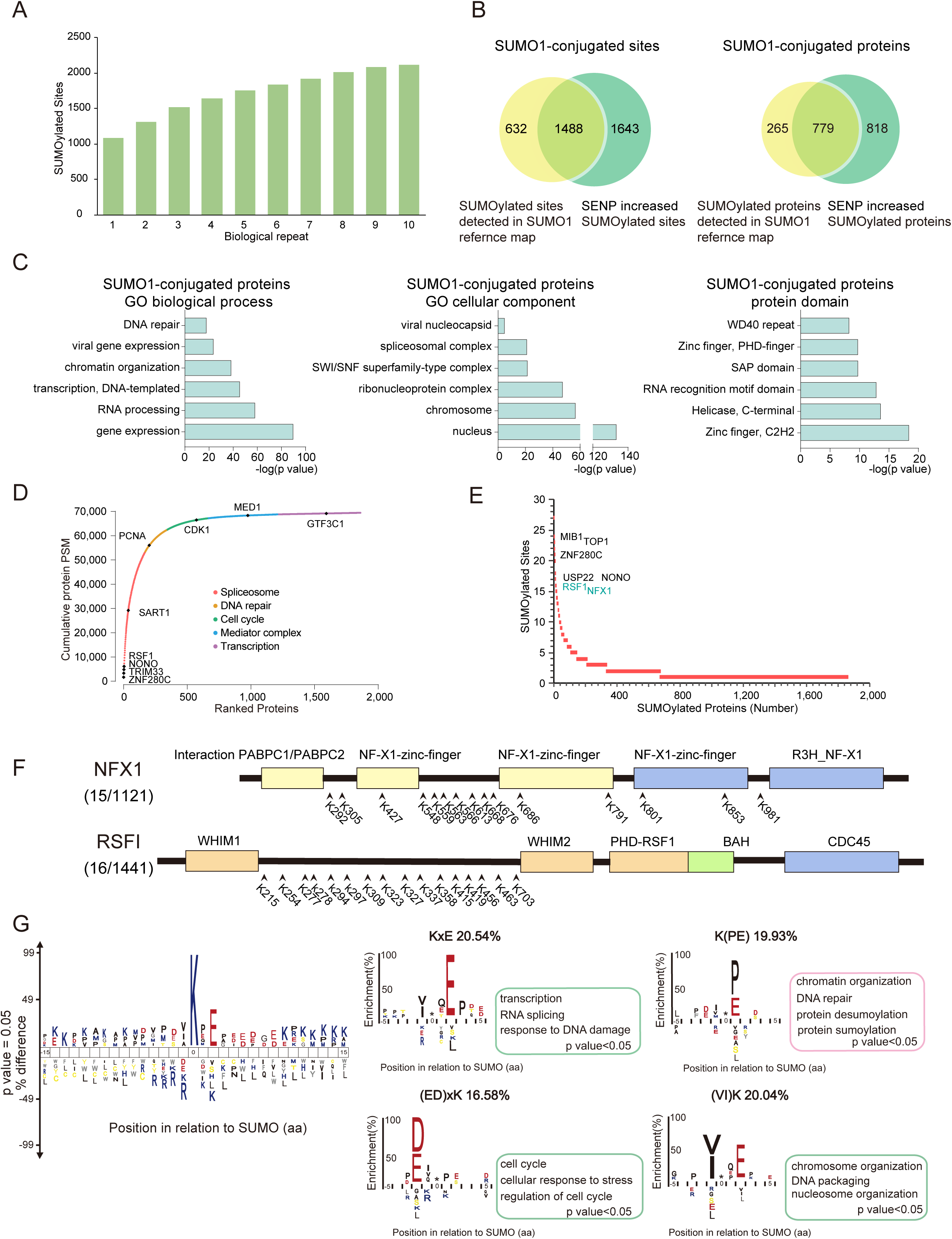
Bioinformatics analysis of SUMO1-conjugated proteins and sites. (A) Bar plot shows the number of detected SUMO1 sites in each of ten biological repeats of the SUMO1 reference map. (B) Venn plot represented SUMO1-modified sites and proteins in SUMO1 reference map and dynamic map. (C) GO enrichment analysis of potential SUMO1-modified proteins. The bar plot indicated the top terms of significantly enriched biological processes, cellular components and protein-domain families for potential SUMO targets. (D) Overview of the SUMOylated peptides identified with their correspondent SUMOylated proteins. The statistical analysis of SUMOylated peptides PSM with the top terms of significantly enriched biological processes were performed. (E) The number of SUMO1 modified sites identified per protein were represented. (F) Graphical representation of the location of SUMO1-modified sites for proteins NFX1 and RSF1. (G) IceLogo representations of SUMOylation sites identification. The amino acids surrounding the SUMO-conjugated lysine under control conditions were analyzed (G, left panel). SubLogo representations of consensus motif in SUMOylation sites. All amino acid changes were significant with P <0.05 by two-tailed Student’s t test. Asterisks symbolled the position of the SUMOylated lysine residue.

Surveying the SUMO1 dynamic map revealed that each SENP members may impact specific signaling pathways. As indicated in Supplementary Figure 3, SENP1 dependent proteins are enriched in response to extracellular stimulus, SENP2-dependent regulation of autophagy, SENP3 in the apoptotic signaling pathway, while SENP5 in viral transcription, SENP6 in DNA repair and SENP7 in the DNA repair and phosphorus metabolic process (**Supplementary Figure 3a, b**).

Collectively, we identified 3,852 sites, and by excluding 89 detected diGly sites identified in His_10_-SUMO1^ΔGG^ cells or in His_10_-SUMO1^WT^ cells, we identified 3,763 high confident SUMO1-conjugated sites representing 1,861 proteins in total (**Figure 1f**). The dataset covered a large number of well-characterized SUMO1-target proteins, including RANGAP1, RANBP2, PCNA, SAFB2, topoisomerases 1, 2α, RNF111, and PML. To evaluate our SUMO1 modified dataset, we compared our data with pervious published SUMO1 data set^23^. For the 295 reported SUMO1 sites 187 sites (63%) were identified in our dataset, and 2,890 novel sites were exclusively identified in our dataset, which indicated our data expanded the coverage of SUMO1-modified peptides with an order of magnitude. (**Supplementary Figure 3c, d, Supplementary Table 1**).

### Bioinformatics analysis of the SUMO1 proteome

We then investigated the basic bioinformatics features of the SUMO1^T95R^ modified reference and dynamic map (the dataset with 3,763 sites, 1,861 proteins). We performed Gene Ontology (GO) analysis (**Supplementary Material and Methods**) on the 1,862 SUMOylated proteins, and the analysis revealed a significant enrichment of SUMO1^T95R^ proteome in the biological processes of gene expression, RNA processing, transcription and DNA repair. The subcellular localization was enriched in nucleus, consistent with previous analysis. Protein with structural domains, such as Zinc finger, RNA recognition motif, SAP domains and WD40 repeat, were identified in large numbers (**Figure 2c, Supplementary Table 2**), indicating the involvement of SUMO1 in the gene transcriptional regulations.

We calculated a cumulative curve for the SUMO1^T95R^ modified site identifications according to peptide spectrum matches (PSM). As shown in the **Figure 2d**, the most abundant SUMOylated proteins, including ZNF280C, TRIM33, NONO, RSF1, account for 8.85% of the PSM of SUMO1 modified peptides. The functional annotations of spliceosome, cell cycle DNA repair, Mediator Complex, and Transcription regulation and cell cycle ranked as the dominant biological processes of SUMOylated proteins (**Figure 2d**). We found MIB1, TOP1, ZNF280C contains most SUMOylated sites (**Figure 2e**). To be more specific, NONO (Non-POU Domain Containing, Octamer-Binding), which is a DNA and RNA binding protein involved in variety of nuclear processes, contained a large number of SUMO1 sites of 16. The E3 ligase MIB1 of the NOTCH pathway and the ubiquitin specific peptidase 22 (USP22) contained 24 and 17 SUMO1 sites, respectively, indicating a potential link between SUMOylation and ubiquitination (**Supplementary Figure 4a**). NFX1 and RSF1, which are involved in the virus infection and inflammatory response, each had more than 10 SUMO1^T95R^ modified sites, suggesting a potential role of SUMO1 in the immune response (**Figure 2f**).

The classical consensus motif of SUMOylation is well established. The commonly observed canonical consensus is ψKxE, where ψ represents hydrophobic residue and x represents any amino acid. We found that approximately 20.54% of SUMO1^T95R^ modified peptides matched the core KxE motif and 16.58% of them matched the inverted (ED)xK motif. At the position-1 of SUMOylated lysine, the preference of large hydrophobic residues including Val and Ile and, showed a 20.04% enrichment over background frequency (**Figure 2g**). Additionally, for the site detected at the position +1 of SUMOylated lysine, Pro and Glu were presented about 19.93% (**Figure 2g**). The GO analysis has revealed that the preference of different motifs is implicated in different biological processes. For instance, the KxE motif is frequently found in proteins involved in transcription and RNA splicing, and the motif K(PE) was enriched in proteins in SUMOylation and deSUMOylation system. Moreover, the inverted (ED)xK was found in cell cycle proteins and the (VI)K was enriched in proteins responsible for chromosome organization. (**Figure 2g**).

### Differential deSUMOylation patterns of the SENP members

We then focused on analyzing deSUMOylated proteins that were dependent on SENPs. SENP-specific deSUMOylation proteins was defined as those whose abundance were at least 2-fold lower in the SUMO1 dynamic map than in the SUMO1 reference map (**Supplementary Materials and Methods**). We found that 378 proteins that were deSUMOylated in a SENP-dependent manner, ranging from 239 proteins for SENP1, to 48 proteins for SENP7 (**Figure 3a**, and **Supplementary Table 3**). As a validation, we confirmed the deSUMOylation specificity of SNIP1 and ADAR1, which were potential substrates of SENP2 and SENP1, respectively (**Figure 3b**). We combined the deSUMOylated proteins of all six SENP members together to profile the sequence motif (**Supplementary Figure 5a**) and bioinformatics features of the SENP specific proteins and found no obvious difference from the SUMOylation system (**Supplementary Figure 5b**).

**Figure 3.**
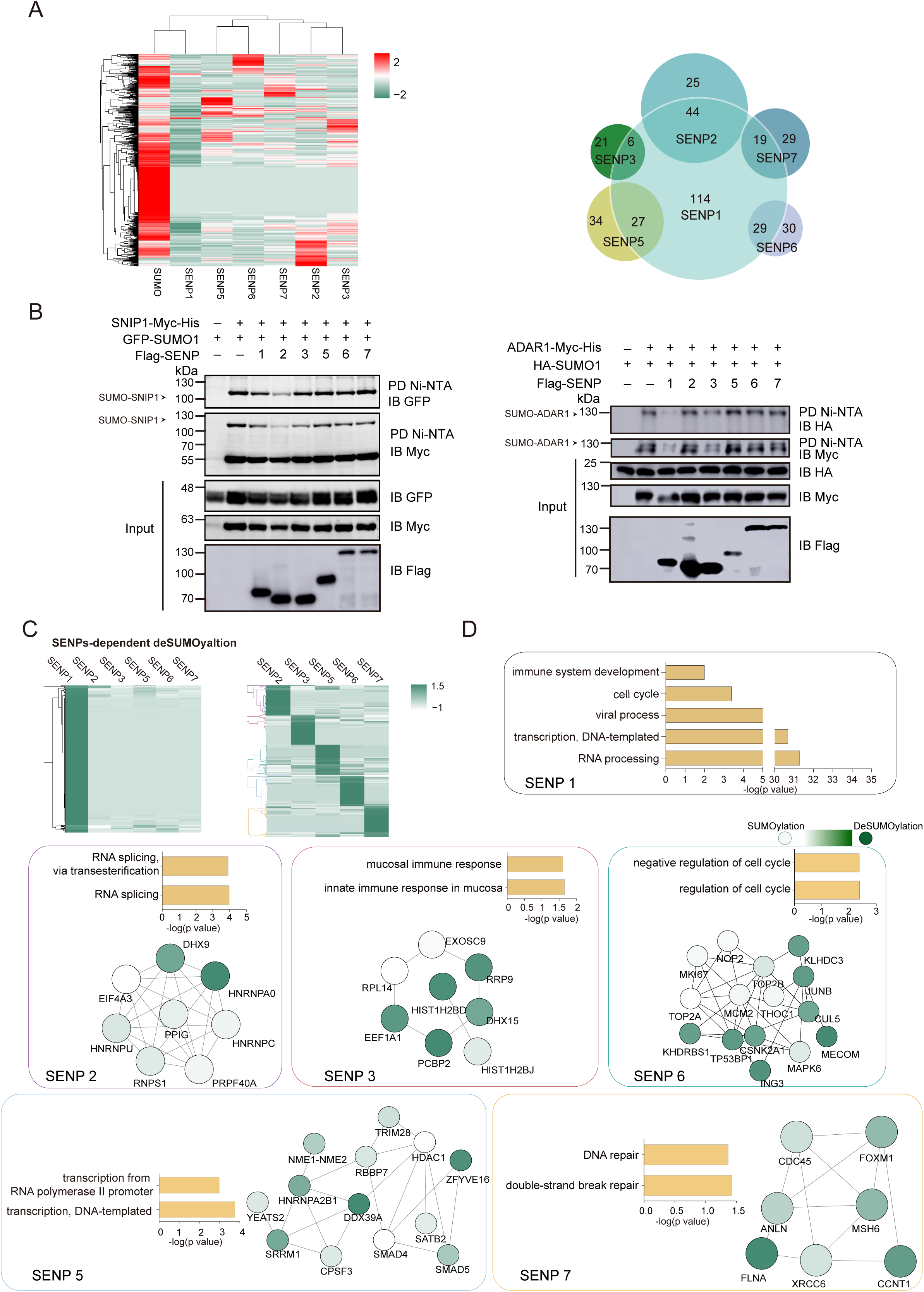
SENP family members modulate dynamic SUMOylation. (A) The cluster heatmap indicated the changes abundance of SUMOylated proteins in SUMO1 reference map and SUMO1 dynamic map. The color bar indicated normalized z-scored FOT (A, left panel). Venn plot shows the number proteins specific deSUMOylated by each SENP member (A, right panel). (B) Immunoblot validated SNIP1 and ADAR1 were specifically deSUMOylated by SENP 2 and SENP1. HEK293T cells were transfected with indicated plasmids, Cell lysates were subjected to Ni-NTA pulldown (Ni-PD), and the deSUMOylated forms of candidates were assayed by indicated antibodies. (C) Clustering heatmap represented the changes of abundance of SUMOylated proteins in SUMO1 dynamic map. The color bar indicated normalized z-scored FOT. (D) The bar plot represented the bioprocess enriched by the proteins deSUMOylated by each SENP family members. The STRING network indicated the interaction among the bioprocess specific proteins. Color bar indicated the deSUMOylated level of each protein.

### SENPs may regulate different bioprocesses revealed by differential SUMO1-SENP modification patterns

As shown in figure 3c, SENP family members can be classified by their deSUMOylation ability. The GO analysis on SENP-specific deSUMOylated proteins were then performed and the main GO biological processes enriched by them (p value<0.05) were compared. The GO biological process analysis indicated that SENP1, as the dominant SENP for SUMO1 sites deSUMOylation, is involved in various biological processes, including transcription, RNA processing, cell cycle, immune system, among others. Importantly, other SENP members display apparent functional preference in different biological processes. For example, the SENP2 participates preferentially in RNA splicing; SENP3 deSUMOylates several proteins involved in the defense response to virus, such as PCBP2, DHX15, EXOSC9. SENP5 is involved in the transcription, and SENP6-dependent deSUMOylation is predominantly involved in the cell cycle progression. Finally, SENP7 is involved in the DNA repair and associated with the proteins in Ub system such as PSMD1, suggesting an involvement of SENP7 in proteasome-mediated protein catabolic process^27^. The GO terms and protein network of the GO bioprocesses dominated with different SENP members was summarized in **Figure 3d**.

Previous analyses indicated that the SENPs deSUMOylate proteins in the cellular pathways of RNA splicing, cell cycle and DNA repair. We investigated the dynamics of SUMO1 modification sites and their dependence on SENPs for proteins involved in these three major pathways.

In the spliceosome, we found that multiple components of U1, U2, U4, U5 and U6 of the major spliceosome were SUMOylated, and they are mainly deSUMOylated in a SENP1- or SENP2-dependent manner (**Figure 4a**). Moreover, we detected many SUMOylation and deSUMOylation events on cell cycle related protein APC/C complex (Anaphase-promoting complex). The site K683 of ANAPC5, site K490 of CDC20 were SUMOylated and deSUMOylated majorly by SENP2 and SENP3. Since APC/C complex is an E3 ubiquitin ligase, we then tried to observe if other proteins participated in ubiquitin mediated proteolysis were deSUMOylated by SENPs. As expected, we found that proteins belonged to E1, E2 and E3 were SUMOylated and deSUMOylated by various SENP members (**Figure 4b**).

**Figure 4.**
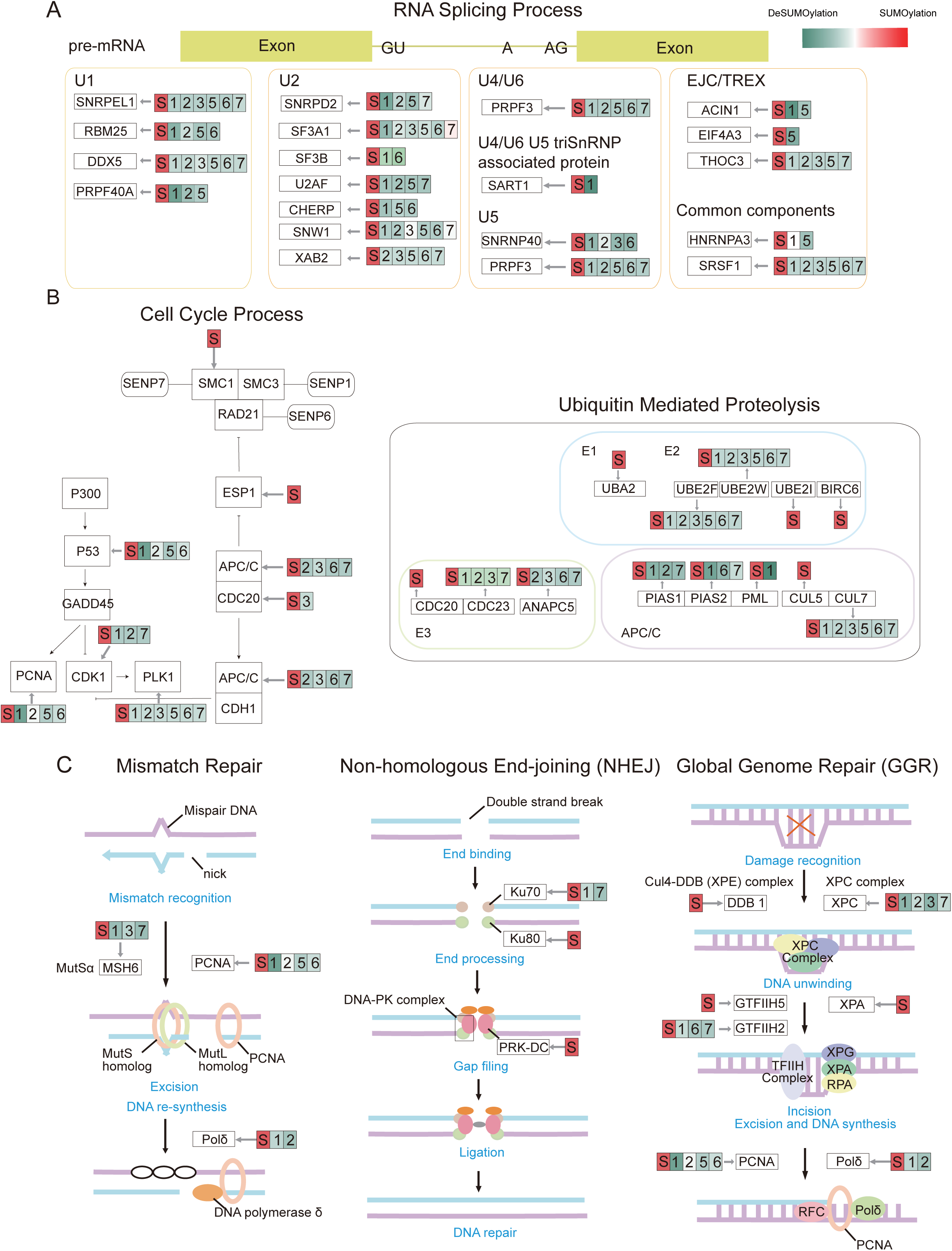
The dynamics of SUMOylation and deSUMOylation throughout cellular pathways. (A) Graphic depiction of dynamically SUMOylated and deSUMOylated proteins participate in RNA splicing mechanism. (B) Systematic overview of dynamically SUMOylated and deSUMOylated proteins participate in cell cycle process. (C) DNA repair related proteins are SUMOylated and deconjugated with SUMO1 by SENP1/3/7 under different conditions. The red S indicated proteins which were SUMOylated. The green numbers indicated proteins which are deSUMOylated by different SENP members.

A pervious study demonstrated that SUMOylation plays an important role in response to DNA damage ^28^. We observed that several core-regulators involved in three different DNA repair processes including mismatch repair, non-homologous end-joining and nucleotide excision repair were SUMOylated and deSUMOylated (**Figure 4c**).

### SENP3 participates in the immune response

The analysis of SENP specific deSUMOylation pattern has revealed that the SENP3 may function in immune response to viral stimulation. We next examined whether SENP3 can mediate RNA agonist induced immune activation. HEK293 cells over-expressing SENPs was treated with poly (I:C) and the expression of anti-viral-responsive genes (*IFNβ, ISG56, RANTES* and *IL8*) were measured by real time PCR (**Figure 5a**). Exogenous expression of SENP3 drastically increased the anti-viral genes expression in response to poly (I:C) challenge than that if other SENP members. In addition, SENP3-C532S mutant, which is the catalytically inactive mutant of SENP3, could not promote the induction of anti-viral genes expression during poly (I:C) infection (**Supplementary Figure 6a**). Furthermore, SENP3 siRNA (SENP3 siRNA 1416) was employed to confirm the regulatory role of SENP3 in response to the poly (I:C) stimuli. Importantly, silencing of SENP3 apparently down-regulated the expression of anti-viral genes (**Supplementary Figure 6b**). As the poly (I:C) mimics the RNA virus, vesicular stomatitis virus (VSV) infection was utilized to further confirm the function of SENP3 in the immune response. We challenged the HEK293 cells directly with VSV-GFP for 8h, or knocked down SENP3 with siRNA and treated the cells with VSV-GFP. SENP3 knockdown resulted in a significant increase in virus titer compared to non-transfected cells (**Figure 5b**), demonstrating that SENP3 plays important roles in anti-viral immune response.

**Figure 5.**
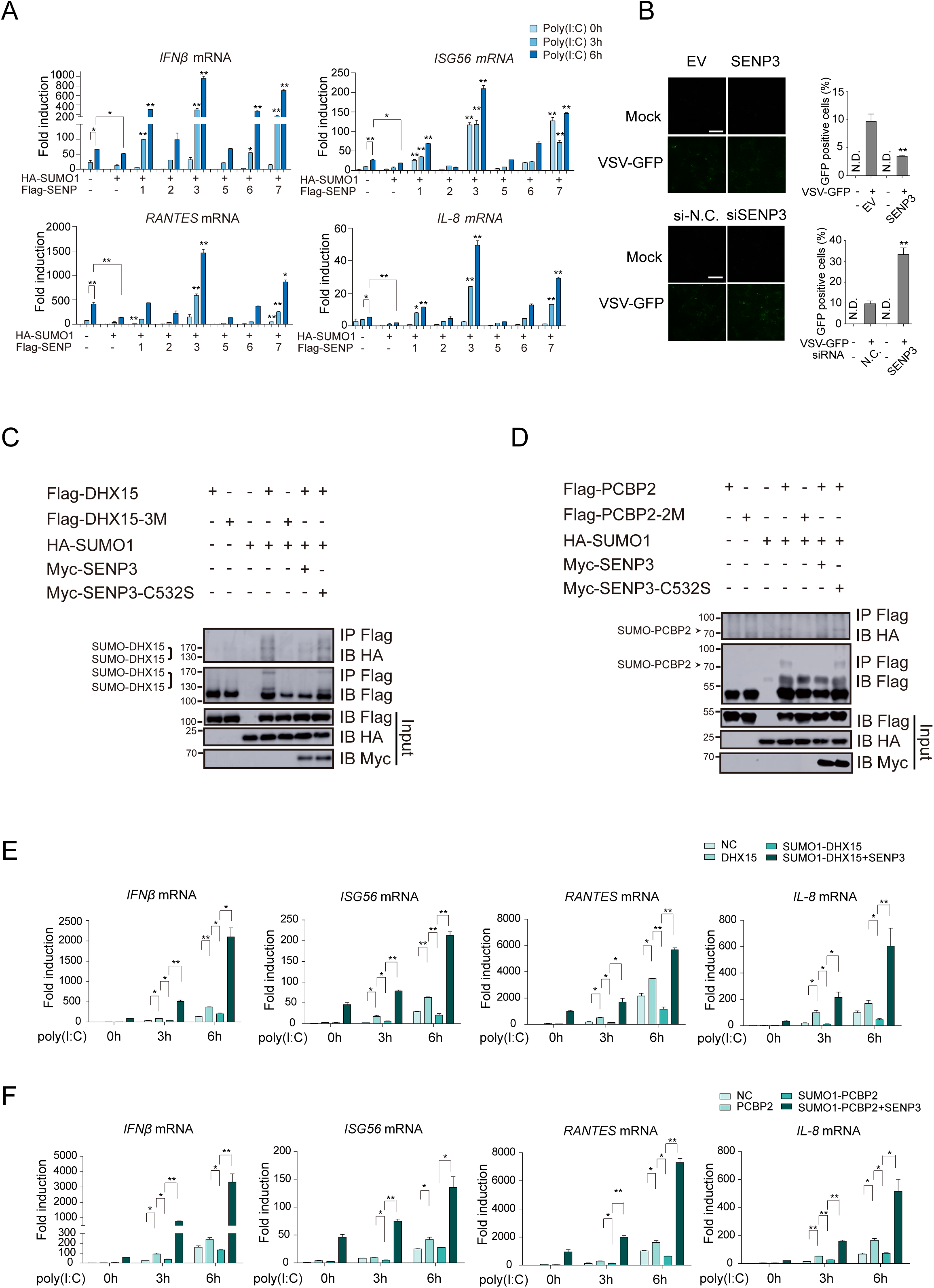
The deSUMOylation function of SENP3 plays a key role in RNA-virus induced signaling pathway. (A) HEK293 cells transfected with the indicated plasmids were stimulated with poly (I:C). Induction of *IFNβ, ISG56, RANTES* and *IL-8* are measured by quantitative PCR. 2μg/ml poly (I:C) was delivered to the cells by Lipofectamine 2000. Graphs showed the mean ± s.d. and data shown were representative of three independent experiments. *p < 0.05; **p < 0.01 (two-tailed t-test). (B) VSV-GFP replication in HEK293 cells transfected with indicated plasmids or siRNA were visualized by fluorescence microscopy. The images were captured with a ×60 objective. Data are representative of triple biological repeats. EV, empty vector, N.C., nonspecific control. (C, D) SENP3 catalyzed deSUMOylation of DHX15 and PCBP2. Flag-tagged human DHX15 and PCBP2 or their mutants were individually transfected into HEK293T cells with HA-tagged SUMO1. 40h post-transfection, cell lysates were subjected to immunoprecipitation and then immunoblotted with the indicated antibodies. (E, F) Quantitative PCR of *IFNβ, ISG56, RANTES* and *IL8* expression from HEK293 cells transfected with the indicated plasmids after stimulation with 2μg/ml poly (I:C) delivered to the cells by Lipofectamine 2000. Graphs showed the mean ± s.d. and data shown were representative of three independent experiments. *p < 0.05; **p < 0.01 (two-tailed t-test)

### SENP3 catalyzes the deSUMOylation of DHX15 and PCBP2 to regulate the antiviral response

To investigate the mechanism of SENP3 in regulating immune response, we analyzed its deSUMOylation patterns to identify potential substrates that could modulate antiviral process. DHX15 has been reported previously as a novel RNA virus sensor via its binding to poly (I:C) to activate immune response^29^, and PCBP2 has also been reported as a negative regulator in MAVS-mediated antiviral signaling through its SUMOylation that causes MAVS degradation^30,31^. In our study, both DHX15 and PCBP2 were dependently deSUMOylated by SENP3 (**Figure 3e, and Supplementary table 3**). To further validate the deSUMOylation, co-transfection experiments were executed (**Figure 5c, 5d**). Flag-tagged DHX15 and PCBP2 were co-expressed with HA-tagged SUMO1 and the cell lysate was co-immunoprecipitated with the anti-Flag antibody. Both DHX15 and PCBP2 were SUMOylated in the presence of SUMO1. Notably, the modifications were dramatically decreased by the overexpression of SENP3. Additionally, knocking down endogenous SENP3 could increase the basal SUMOylation levels of Flag-DHX15 and Flag-PCBP2 (**Supplementary Figure 6c, Supplementary Figure 6d**), suggesting that SENP3 catalyzes deSUMOylation of DHX15 and PCBP2.

To identify the potential regulatory SUMO1 site, we examined the SUMOylation sites on DHX15 and PCBP2 individually depend on SUMO1 reference map and dynamic map. Lysine 17, 18 and 754 of DHX15, and Lysine 115 and 119 of PCBP2 were identified to be indispensable for SUMOylation. We generated DHX15-3M mutant (K17R, K18R and K754R) and PCBP2-2M mutant (K115R and K119R) respectively to investigate the biochemical and functional consequences of these mutations. As shown in **Figure 5c and 5d**, the SUMOylation signals of both mutants were largely abolished (**Figure 5c, 5d, compare lane 5 to lane 4**). We next determined the effect of the SENP3-dependent deSUMOylation of DHX15 and PCBP2 in poly (I:C)-induced immune response. As shown in **Figure 5e, 5f**, over-expression of DHX15 or PCBP2 resulted a moderate increase in anti-viral-responsive genes expression (*IFNβ, ISG56, RANTES* and *IL8*) in response to poly (I:C) stimulation in HEK293 cells; however, the increased response was dampened by co-expression SUMO1, suggesting the SUMOylation plays an inhibitory role in the cytosolic RNA-triggered anti-viral genes expression. Importantly, ectopic-co-expression of SENP3 could not only reversed the SUMO1-induced dampening, it promoted even higher levels of IFN transcription (**Figure 5e, 5f**), Furthermore, the deSUMOylation mutant of DHX15 (DHX15-3M) and PCBP2 (PCBP2-2M) displayed similar immune response effect on the production of anti-viral genes (*IFNβ, ISG56, RANTES* and *IL8*), and these promoting effects could not be diminished by the overexpression of SUMO1, consistent with a role for SENP3-dependent deSUMOylation of DHX15/PCBP2 in promoting immune response (**Supplementary Figure 6e, Supplementary Figure 6f**). Collectively, these results suggested that SENP3 plays a role in anti-viral response and the deSUMOylation activity of SENP3 is critical in this process.

### The differential interactome of SENP3 that correlates with the antiviral immune response

To provide additional evidence for SENP3 in the anti-viral signaling pathway, the immunoprecipitation coupled mass spectrum (IP-MS) of SENP3 was performed to identify proteins in the RNA virus-induced immune response. HEK293 cells were transfected with Flag-tagged SENP3 and incubated with or without poly (I:C) for 6h (EV, SENP3, EV poly I:C, SENP3 poly I:C). Whole cell lysates were prepared and subjected to anti-Flag immunoprecipitation. Co-precipitated proteins were separated by SDS-PAGE and analyzed by LC-MS (**Figure 6a**). We compared the SENP3 IP-MS between EV and SENP3 transfected cells and summarized the potential interaction network of SENP3 with or without poly (I:C) stimulation (**Figure 6b**). SENP3 seemed to interact with several key proteins in immune response. For example, binding to the kinase EIF2AK2/PKR suggested a connection of SENP3 with the IFN-induced signaling pathway. In addition, DDX60, TAK1 and IFIT5 were further increased association with SENP3 in response to poly (I:C) stimulation (**Figure 6c**). The GO/pathway analysis of poly (I:C) induced SENP3 interacting proteins showed an enrichment of GO terms in RNA splicing, IκB phosphorylation and RNA processing, suggesting a correlation between SENP3 and the NF-κB signaling pathway under poly (I:C) administration (**Figure 6d**). We assumed that SENP3 modulates immune responses to RNA virus infection in the model presented in **Figure 6e**, bridging the SUMO-SENP system to the immunity.

**Figure 6.**
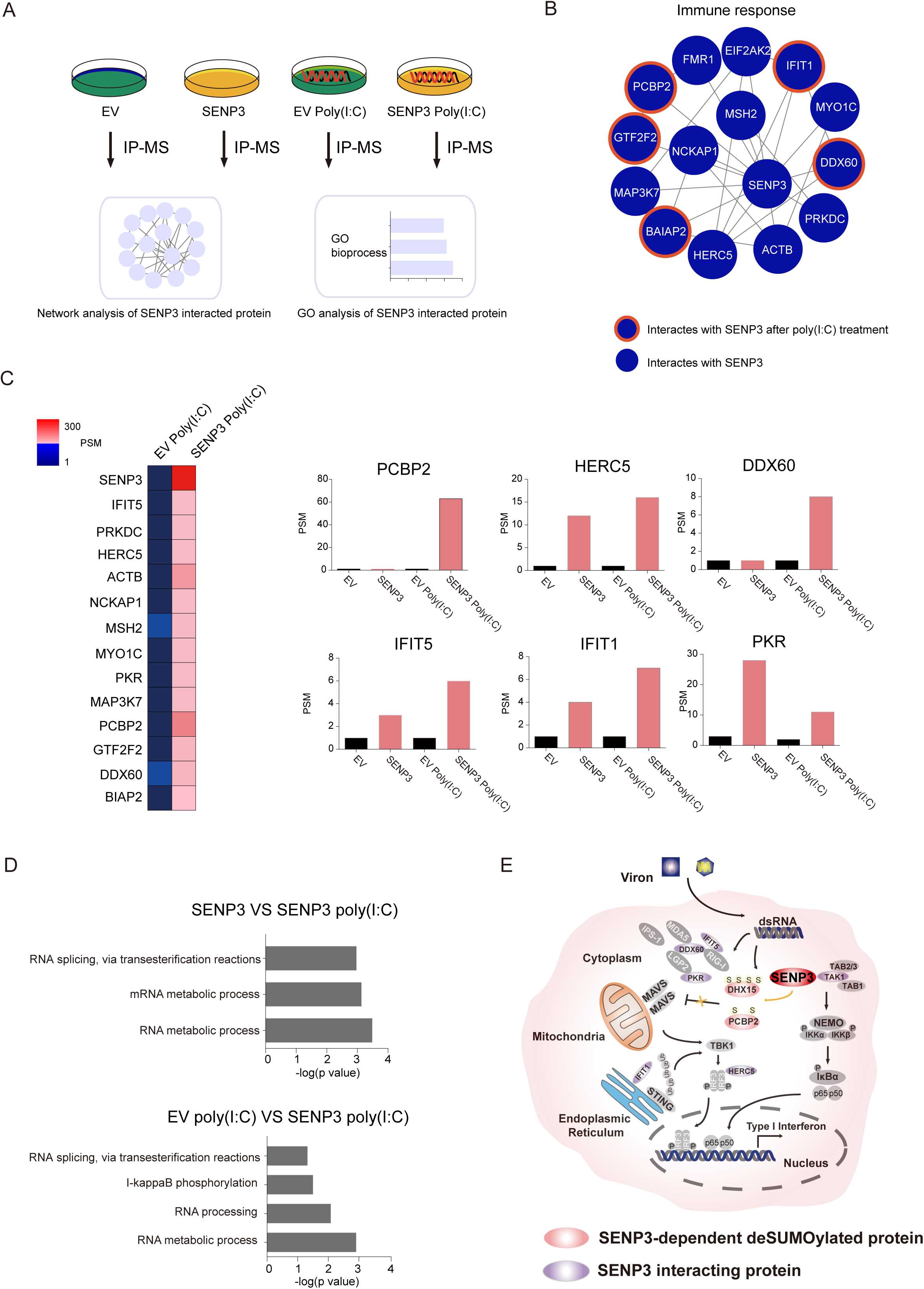
SENP3 promotes antiviral immunity during poly (I:C) stimulation in HEK293 cells. (A) Schematic representations of SENP3 interactome identification during poly (I:C) stimulation in HEK293 cells. EV and Flag-tagged SENP3 were transfected into HEK293 cells and poly (I:C) stimulation was performed 6hr before cell collection. The lysates were subjected to immunoprecipitation and then in gel digestion for LC-MS/MS. Three biological replicates were performed for data analysis. (B) The interaction network of SENP3 in HEK293 cells, with or without poly (I:C) treatment. (C) Increased association of SENP3 interacted inflammatory proteins under poly (I:C) stimulation and the expression profiles were represented. (D) The GO/pathway enrichment analysis of SENP3 interactome under poly (I:C) stimulation. (E) Potential mechanism of SENP3 in regulating RNA-virus induced immune signaling pathway.

## Discussion

The reversible post-translational modifications (PTMs) via small molecules and proteins enable the precise control of protein activations. The deep coverage of SUMOylation will provide new insights into the regulations of biological processes orchestrated by SUMO system. The strategy employed in this study combined peptide immunopurification protocols with mass spectrometry analysis to obtain proteome-wide identification of SUMOylation sites^22,32^. Recently, the SUMOylation patterns of SUMO2 and SUMO3 were provided with deep coverage. In contrast, the largest dataset for SUMO1-modified sites reported by Impens et al. contains only 295 sites^23^. Here, we applied an affinity purification of SUMO1^T95R^-modified proteins combined with anti K-ε-GG immunoprecipitation to achieve the identification of almost 4,000 SUMO1-conjugated sites on over 1,800 proteins, representing the deepest SUMO1-modified proteome. Previous studies indicated that SUMOylation primarily occurs on the classical forward consensus motif ψKxE/D or the inverted consensus motif E/DxK^23,33^. Our study has confirmed that majority of SUMO1-modified sites occur on the typical consensus motifs. Importantly, with the expanded the coverage of SUMO1-modified peptides (>10 times), we have not only observed over 30% SUMOylated peptides possess consensus adherence under standard growth conditions, but also identified a novel sub-motif K(PE) with Pro and Glu located +1 position of the SUMOylated lysine. The GO analysis has revealed the enrichment of chromatin organization, DNA repair and SUMOylation and deSUMOylation processes in the K(PE) motif, suggesting a functional enrichment for this motif.

We initially generated His_10_-SUMO1^T95K^ stable cell line and tried the approach described by Tammsalu et al^32^, to eliminate the endogenous modification by ubiquitin and ubiquitin like modifiers. However, the commercial K-ε-GG antibody was not suitable to enrich peptides generated by Lys-C digestion, and the relatively small proportion of diGly remnant attached to the target lysines was identifiable. For this reason, we choose His_10_-SUMO1^T95R^ mutant cell line coupled with trypsin instant to construct SUMO1 reference map.

Indeed, His_10_-SUMO1^T95R^ has its limitation that peptides modified by Ub and Ubl will leave the same diGly remnants after trypsin digestion, thus to filter SUMO1 modified peptides against peptides modified by Ub and by other Ubls, we set His_10_-SUMO1^ΔGG^ mutant cell line and His_10_-SUMO1^WT^ mutant cell line as background control and exclude diGly sites identified in those cells. By carefully comparing the diGly modified peptides detected in His_10_-SUMO1^T95R^ cell line and His_10_-SUMO1^ΔGG^ mutant cell line and His_10_-SUMO1^WT^ mutant cell line, we found the abundances of diGly modified peptides detected in His_10_-SUMO1^T95R^ cell line are higher than peptides detected in His_10_-SUMO1^ΔGG^ mutant cell line or in His_10_-SUMO1^WT^ mutant cell line, indicating we may rule out some potential SUMO1 modified sites as well, which conforms the strictness of our strategy.

The SENP family is constituted of six members, SENP1, 2, 3 and SENP5, 6, 7. SENP1 is known as the major deconjugase of SUMO1 in the SENP family. The subfamilies were classified based on the sequence homology and substrate specificity. SENP1 and SENP2 consist of the first subfamily and shared deconjugation functions through SUMO community ^8^; SENP3 and SENP5, which belong to the second subfamily, are enriched in the nucleoli and involved in ribosome biogenesis ^8,14^. The third subfamily consisting of SENP6 and SENP7 display higher specificity for SUMO2/3 than SUMO1 ^8^. However, as the previous studies predominantly focused on the investigations of SENPs individually, functional features of different SENP members could not be compared. The identification of substrates of SENP family is critical to understanding the dynamic of SUMOylation and its modulation to respond external and internal signals. In our study, due to the inherent low abundance of SENPs’ expression in HEK293T cells, SENP family members were over-expressed individually in SUMO1-expressing cells and potential substrates of each SENP family members were identified by differentially deSUMOylated sites. As illustrated by our dataset in SUMO1-modified proteome and previous studies, SENP1 is the ubiquitous SUMO1 deconjugase ^6^. SENP2 facilitates RNA splicing process and promotes nuclear activities ^13,34^. SENP5 mediates transcription and is essential for cell proliferation ^35,36^. SENP6 and SENP7 regulate cell cycle and DNA repair respectively^17,37,38^.

The host innate immune system monitors invading cytosolic microbial RNAs and initiates antiviral responses. Several sensors have been characterized to recognize viral RNA and transmit the activation signal to the mitochondria-resident protein MAVS, which activates the TBK1 kinase and IRF3 transcriptional factor to produce type-I interferons (IFNs) and inflammatory cytokines ^39^. DHX15, a member of DEXD/H-box family helicase, was confirmed to be a novel RNA sensor which was required to bind poly (I:C) and interacts with MAVS to trigger IRF3 activation and NF-κB and MAPK signaling ^29,40^. By analyzing SUMO1 dynamic map in this study, we speculated that DHX15 is a potential substrate of SENP3. DHX15 could be modified by SUMO1 and the modification sites were mapped onto lysine residues 17, 18 and 754. SUMOylation of DHX15 markedly attenuated the RNA-triggered expression of IFNs and ISGs, while the effect could be rescued by exogenously expressed SENP3. We also identified TAK1 (also known as MAP3K7), which phosphorylates and activates IKK complex ^41^, could be recruited by SENP3 to induce NF-κB signaling pathway. As for PCBP2, the host RNA-binding protein PCBP2 (poly (rC) binding protein 2) was reported to be SUMOylated by SUMO2 at lysine 37 in the nucleus and translocate to the cytoplasm ^31^. The SUMOylated PCBP2 associates with MAVS to promote its proteasomal degradation, leading to down-regulation of MAVS signaling cascade and ‘fine tuning’ of antiviral innate immunity ^30^. We found that PCBP2 could also be SUMOylated by SUMO1 at lysine 115 and 119 and SUMOylated PCBP2 obviously suppressed the induction of IFNs and ISGs, indicating that the modifications by SUMO1 and SUMO2 shared comparable functions. Recently, Hu et al reported that RIG-I and MDA5 could be modified with SUMO1 by E3 ligase TRIM38 catalysis and deSUMOylated by SENP2 at the late phase of viral infection ^42^. However, our cell-based purifications failed to detect RIG-I and MDA5 as potential substrates for SUMO1 modification. We speculated that the different cell types utilized in our studies, which may contain different sets of SUMOylated proteins, is a possible reason for the discrepancy. Another possibility was that the study of Hu carried out in the presence of E3 ligase TRIM38 that lead to SUMOylation reaction of RIG-I and MDA5, whereas our study was performed without specific E3 ligases. Interestingly, SENP2 was proved to deSUMOylate RIG-I and MDA5 and promote their K48-linked polyubiquitination and degradation at the late phase of RNA virus infection (8-16h) to avoid sustained activation of RIG-I and MDA5, indicating that SENP2 is a negative regulator in the MAVS signaling cascade. Our study focused on the early phase of RNA virus infection (0-6h) and showed that SENP3 not only deSUMOylates DHX15 and PCBP2 to increase transcription of downstream antiviral genes but also recruits DDX60 and HERC5 to promote RLR-mediated antiviral activities. These results indicate that SENP family members cooperates to synergistically modulate cellular response to pathological conditions, and reveal vital roles of SENP members in innate immune response.

Collectively, this study developed a streamlined strategy that allowed the deep coverage of SUMO1-modified proteome, uncovered the biological preferences of different SENPs by monitoring the dynamic of SUMOylation/deSUMOylation of SENP family. We further demonstrated the involvement of SENP3 in the antiviral immune response and identified potential substrates functioning in the process. The SUMO1-modified proteome and SENP-deSUMOylation network in this study provide a useful resource for understanding and connecting the biological functions with the SUMOylation/deSUMOylation systems.

## Materials & Methods

### Plasmids

Human full-length SENP1, SENP2, SENP3, SENP5, SENP6, SENP7, SNIP1, ADAR1 and PCBP2 cDNA were cloned from human thymus plasmid cDNA library (Clontech) using standard PCR techniques and then subcloned into indicated vectors. DHX15 constructs were kindly provided by Dr. Jiahuai Han (Xiamen University). All point mutants were generated by using a Quickchange XL site-directed mutagenesis method (Stratagene). All constructs were confirmed by sequencing.

### Antibodies and reagents

The SUMO1 antibody and β-actin antibody were purchased from Abcam. The antibodies against hemagglutinin (HA), Myc and ubiquitin (41) were purchased from Abmart. Anti-Flag (M2)-agarose, flag antibody, poly(I:C), CHX and MG132 were obtained from Sigma. PTMScan Ubiquitin Remnant Motif (K-ε-GG) Kit were purchased from Cell Signaling Technology (CST#5562).

### Construction of HEK293T cells stably expressing His_10_- SUMO1^T95R^, His_10_- SUMO1^T95K^, His_10_- SUMO1^WT^, His_10_- SUMO1^ΔGG^

pCDH-CMV-MCS-EF1-Puro-His_10_-SUMO1^T95R^, pCDH-CMV-MCS-EF1-Puro-His_10_-SUMO1^T95K^, pCDH-CMV-MCS-EF1-Puro-His_10_-SUMO1^WT^, pCDH-CMV-MCS-EF1-Puro-His_10_-SUMO1^ΔGG^ were constructed by cloning a PCR-generated 10His-SUMO1 fusion into the EcoRI and BamHI sites of the plasmid vector pCDH-CMV-MCS-EF1-Puro. The SUMO1^T95R^, SUMO1^T95K^, SUMO1^ΔGG^ mutation were introduced into pCDH-CMV-MCS-EF1-Puro-His_10_-SUMO1 by site-directed mutagenesis. The SUMO1 coding plasmids were verified by sequencing. HEK293T cells stably expressing His_10_-SUMO1^T95R^, His_10_-SUMO1^ΔGG^, His_10_-SUMO1^T95K^, His_10_-SUMO1^WT^ were generated through lentiviral infection with a virus encoding pCDH-CMV-MCS-EF1-Puro-His_10_-SUMO1^T95R^, pCDH-CMV-MCS-EF1-Puro-His_10_-SUMO1^T95K^, pCDH-CMV-MCS-EF1-Puro-His_10_-SUMO1^WT^, pCDH-CMV-MCS-EF1-Puro-His_10_-SUMO1^ΔGG^ which were packaged for 48h. The selected cells were cultured in the presence of puromycin at 3μg/ml.

### Cell culture and transfection

HEK293T (ATCC) and HEK293 (ATCC) cells were cultured in DMEM (Invitrogen) plus 10% FBS (Gibco), supplemented with 1% penicillin-streptomycin (Invitrogen). Cells (∼3×10^7^) were cultured for 48h, harvested by centrifugation and washed with cold 1×PBS. Lipofectamine 2000 (Invitrogen) was used for transient transfection of HEK293T and HEK293 Cells according to the manufacturer’s instructions

### RNA interference and manipulation of virus

The siRNA duplexes targeting SENP3 was chemically synthesized by Gene-Pharma. The sequences of siRNAs are shown as follows: *human SENP3*, 5’-GCU UCC GAG UGG CUU AUA ATT-3’; The nonspecific siRNA (si-N.C.), 5’-UUC UCC GAA CGU GUC ACG UTT-3’. VSV-GFP was kindly provided by Dr. Feng Qian (Fudan University). Viral infection was performed when 80% cell confluence was reached and VSV-GFP was added to the media at MOI=1.

### Protein extraction and nickle affinity purification

Cell pellets were lysed in ten pellet volumes of guanidine lysis buffer (6 M guanidine-HCl, 100 mM sodium phosphate, 10 mM Tris-HCl, 20mM imidazole and 5mM β-mercaptoethanol, buffered at pH 8.0) supplemented with 10mM PMSF (Sigma). Lysates were subjected to short pulse (5 s per 5 mL lysate, total sonication time of 15 s) of sonication with sonicator at a power of 30W. Insoluble particles were removed by centrifugation for 15000g, 15min. Protein concentration was determined by Bradford assay (TAKARA). Nickel affinity purification of 10His-SUMO1^T95R^ conjugates was carried out with Ni-NTA agarose beads (Qiagen). 25μl beads were prepared per 1ml lysate by washing for three times with guanidine lysis buffer and added to the cell lysate for mixing overnight at 4□. After incubation, beads were washed with 10 bead volumes of the following wash buffers in order: guanidine lysis buffer, wash buffer pH 8.0 (8 M urea, 100 mM sodium phosphate buffer pH 8.0, 10 mM Tris-HCl, 10 mM imidazole, 5mM β-mercaptoethanol), wash buffer pH 6.3 (8 M urea, 100 mM sodium phosphate buffer pH 6.3, 10 mM Tris-HCl, 10 mM imidazole, 5 mM β-mercaptoethanol) and again with wash buffer pH 8.0. Proteins were eluted in five sequential steps with 2 bead volumes of elution buffer (8 M urea, 100 mM sodium phosphate buffer pH 8.0, 10 mM Tris-HCl, 200 mM imidazole, 5 mM β-mercaptoethanol).

### Filter aided sample preparation and protein digestion

Digestion of 10His-SUMO1^T95R^ proteins was performed on 10 kDa-cutoff spin filters (Sartorius) according to a published protocol. Samples were concentrated on filters and treated with 1mM dithiothreitol (DTT) in the dark for 30 minutes. Next, chloroacetamide was added to a final concentration of 5mM to incubate with samples for 30min at room temperature. After alkylation, 5mM DTT was supplemented with samples for 30min at room temperature. Samples were then washed twice with urea wash buffer (250μl of 8M urea, 100mM Tris pH 7.5), three times with 250 μl of 50mM ammonium bicarbonate (ABC) buffer and digested for 16 hours with sequencing-grade modified trypsin (Promega) in 200 μl ABC buffer at 37 °C (enzyme to protein ratio 1:50). Sample were collected and the filters were washed twice with 150 μl of HPLC water to increase the yield of peptides. Final concentration of the peptides was dried in a Savant SpeedVac and stored at −80 °C.

### Immunopurification enrichment of diGly-Lys containing peptides

Anti-K-ε-GG conjugated to protein A beads (4μl of beads; PTMScan, Cell Signaling Technology) was washed three times with 1 ml of 1ml 1×PBS, centrifuged at 2000g, 30s after each wash. Then the beads were resuspended in 40μl PBS of each sample. Enrichment of diGly-Lys containing peptides with anti-K-ε-GG was performed according to the manufacturer’s instructions. Anti-K-ε-GG (25μg) cross-linked to protein A beads (4μl) was added to peptide mixtures (approximately 300μg) dissolved in 1×IAP buffer and incubated at 4 °C for overnight while rotating. Beads were washed twice with 500μl of cold 1×IAP buffer and twice with 500μl of HPLC water. The SUMOylated peptides were eluted three times with 100 μl of 0.15% trifluoroacetic acid (TFA) and dried in a Savant SpeedVac for the following LC-MS/MS identification.

### Immunoprecipitation (IP) assay, immunoblot analysis and IP-MS

For immunoprecipitation assay, HEK 293T cells extracts with overexpression of the indicated plasmids were prepared by using NETN buffer (50 mM Tris-HCl pH 8.0, 100 mM NaCl, 1mM EDTA, 0.5% NP-40) supplemented with 10mM PMSF (Sigma). Lysates were incubated with Anti-Flag (M2)-agarose (Sigma) for 4h to overnight at 4□. The immunoprecipitates were washed four times with the NETN buffer and eluted with SDS loading buffer by boiling for 5 min. Then the immunoprecipitates were subjected to immunoblot analysis.

For immunoblot analysis, the samples were separated by SDS-PAGE. The resolved proteins were then electrically transferred to a PVDF membrane (Millipore). Immunoblotting was probed with indicated antibodies. The protein bands were visualized by using a SuperSignal West Pico chemiluminescence ECL kit (Pierce) and monitored by Amersham Imager 600 (GE).

For immunoprecipitation coupled MS, control and SENP3 over-expressed HEK293 cells were stimulated by poly (I:C) for 6h and lysed in NETN buffer supplemented with 10mM PMSF (Sigma). The whole cell lysates were immunoprecipitated with 10 μl Anti-Flag (M2)-agarose (Sigma) by incubating for 4 h at 4°C. Immunoprecipitates were washed with NETN buffer and eluted with SDS loading buffer for SDS-PAGE. All samples were prepared by in-gel digestions and analyzed by LC-MS/MS.

### Real-time RT-PCR

Total cellular RNA was isolated with TRIzol (Invitrogen) according to the manufacturer’s instructions. Reverse transcription of purified RNA was obtained by RevertAid RT Reverse Transcription Kit (Thermo Fisher Scientific). The quantification of gene transcripts was performed by quantitative PCR (Q-PCR) using SYBR Premix Ex Taq (TAKARA). All values were normalized to the level of β-actin mRNA. The primers used were listed as follows. *β-actin*, sense (5’-AAAGACCTGTACGCCAACAC-3’), antisense (5’-GTCATACTCCTGCTTGCTGAT-3’); *IFN-β*, sense (5’-ATTGCCTCAAGGACAGGATG-3’), antisense (5’-GGCCTTCAGGTAATGCAGAA-3’); *ISG (IFN-stimulated gene) 56*, sense (5’-GCCATTTTCTTTGCTTCCCCTA-3’), antisense (5’-TGCCCTTTTGTAGCCTCCTTG-3’); *RANTES*, sense (5’ TACACCAGTGGCAAGTGCTC-3’), antisense (5’-ACACACTTGGCGGTTCTTTC-3’); *IL-8*, sense (5’-AGGTGCAGTTTTGCCAAGGA-3’), antisense (5’-TTTCTGTGTTGGCGCAGTGT-3’).

### MS Analysis

LC-MS/MS analyses were performed on an Easy-nLC 1000 liquid-chromatography system (Thermo Fisher Scientific) coupled with a Q-Exactive HF through a nano-electrospray ion source (Thermo Fisher Scientific). The peptide mixture was eluted from a 360-μm ID × 2 cm, C18 trap column and separated on a homemade 150 μm I.D. × 12 cm column (C18, 1.9 μm, 120 Å, Dr. Maisch GmbH) with a 75-min linear 5–35% acetonitrile gradient at 600nL/min. Survey scans were acquired after accumulating of 3e6 ions in Orbitrap for m/z 300–1400 using a resolution of 120,000 at m/z 200. The top 20 intense precursor ions were selected for fragmentation in the HCD cell at a normalized collision energy of 27%, and then fragment ions were transferred into the Orbitrap analyzer operating at a resolution of 15,000 at m/z 200. The dynamic exclusion of previously acquired precursor ions was enabled at 18s.

### Data Processing

Raw MS data files were searched against human protein RefSeq database (released 1 July 2013, 27414 proteins). with Protein Discover (Thermo Fish Scientific, version 1.4) using MASCOT^43^ search engine with percolator^44^. The mass tolerance of MS/MS spectra was set to 20 p.p.m., and the tolerance of the product ions was set to 50 mmu. Up to two missed cleavages were allowed for protease digestion, and the minimal required peptide length was set to seven amino acids. Database searches were performed with Trypsin (K/R) specificity and four missed cleavage sites were allowed. Peptides were accepted with a minimum length of 7 amino acids and a maximum size of 4.6 kDa, Acetyl (Protein-N term), oxidation (M), NEM (C) and GlyGly (K) were chosen as variable modifications, carbamidomethylation (C) was chosen as a fixed modification. 2 missed cleavages on trypsin were allowed.

The abundance of protein expression was estimated based on the precursor area under the curve. We then used the fraction of total (FOT) to represent the normalized abundance of each protein.

To define the SENP-specific deSUMOylated proteins we calculated the fold change between the FOT of proteins from SUMO reference map to SUMO dynamic map, and for calculation, the non-value quantities were replaced by the minimal FOT of the whole matrix.

### Definition of SENP specific deSUMOylated protein

We classified SUMO1 modified proteins into 6 SENP specific deSUMOylated groups, based on their deSUMOylated level by different SENP members.

As for SENP1-specific deSUMOylated protein, for one thing, the fold change between SUMOylated proteins’ abundance detected in SUMO1 reference map to proteins’ abundance detected in SENP1’s specific SUMO1 dynamic map must be larger than 2. For another, the abundance of the protein detected in this SENP1’s specific SUMO1 dynamic map must be less than 2 folds of the geomean of the proteins’ abundance detected in other 5 SENP’s specific SUMO1 dynamic map. And since the dominant role of SENP1 in deSUMOylating SUMO1 modification, to equally measure the deSUMOylating ability of the other 5 SENP members, we excluded SENP1 from SUMO1 dynamic map, and defined SENP (2, 3, 5, 6 and 7) specific deSUMOylated proteins as follows: for one thing, the fold change between SUMO1 modified proteins’ abundance detected in SUMO1 reference map to proteins’ abundance of expression detected in this SENP’s specific SUMO1 dynamic map must be larger than 2. For another, the abundance of the protein detected in this SENP’s specific SUMO1 dynamic map must be less than 2 folds of the geomean of the proteins’ abundance detected in other 4 SENP’s specific SUMO1 dynamic map.

### Bioinformatics Analysis

Sequence analysis was performed with Motif-X, pLogo and Icelogo ^45–47^. All the peptides for SUMO1 modified sites were pre-aligned and used for the analysis. GO enrichment analysis for biological process, molecular function, and cell component was performed by Database for Annotation, Visualization and Integrated Discovery DAVID (version 6.8), with default setting. Network analysis was performed STRING database (v10.5), and visualized by Cytoscape.

## Acknowledgments

This work was supported by Ministry of Science and Technology of China (Grant 2017YFA0505102, 2016YFA0502500); National Program on Key Basic Research Project (973 Program, 2014CBA02000); National International Cooperation Grant (2014DFB30010, 2012DFB30080 and CPRIT RP110784); National High-tech R&D Program of China (863 program, 2014AA020201 and 2015AA020108); National Natural Science Foundation of China (Grant 31270822); National institute of health (Illuminating Druggable Genome, Grant U01MH105026) and a grant from the State Key Laboratory of Proteomics (Grant SKLP-YA201401).

## Author Contributions

Conceptualization: C.D., R.G., J.Q., F.C.H; Methodology: R.G., Y.Z.W., X.H.W., L.S., K.L., L.L.C., M.W.L., W.H.S.; Formal Analysis: Y.Z.W., R.G., G.Y.Y.; Investigation: R.G., Y.Z.W., C.D., J.Q., F.C.H; Data Curation: Y.Z.W., X.H.W., G.Y.Y; Writing – Original Draft: C.D., R.G., Y. Z.W., J.Q.; Writing –Review & Editing: C.D., R.G., Y.Z.W., J.Q., Y.W.; Funding Acquisition: C.D., J.Q., F.C.H., Y.W., B.Z.; Supervision: C.D., J.Q., F.C.H..

