## Supplementary Appendix for "Global Screening of Sentrin-Specific Protease Family Substrates in SUMOylation"

**Supplementary Figure 1. Comparison between SUMO1-conjugation sites identification by using $His_{10}$-SUMO1$^{T95R}$ mutant and in $His_{10}$-SUMO1$^{T95R}$ mutant (related to Figure 1).**

(A) Scheme workflow of SUMO1-conjugated sites identification in $His_{10}$-SUMO1$^{T95R}$ mutant cells and in $His_{10}$-SUMO1$^{T95K}$ mutant cells.

(B) The bar plot indicates comparison between the proportion of SUMO1T95R modified peptide in the enriched samples (black) and the proportion of SUMO1T95K modified peptide in the enriched samples (gray).

(C) Bar plot shows the comparison between the total abundance of diGly modified peptides detected in $His_{10}$-SUMO1$^{T95R}$ cells and $His_{10}$-SUMO1$^{T95K}$ cells, before and after K-ε-GG antibody enrichment.

(D) Bar plot shows the comparison between the total number of diGly modified peptides detected in $His_{10}$-SUMO1$^{T95R}$ cells and $His_{10}$-SUMO1$^{T95K}$ cells, before and after K-ε-GG antibody enrichment.

(E) Bar plot shows the comparison between the total abundance of peptides detected in $His_{10}$-SUMO1$^{T95R}$ cells and $His_{10}$-SUMO1$^{T95K}$ cells, before and after K-ε-GG antibody enrichment.

(F) Bar plot shows the comparison between the total number of proteins detected in $His_{10}$-SUMO1$^{T95R}$ cells and $His_{10}$-SUMO1$^{T95K}$ cells, before and after K-ε-GG antibody enrichment.

(G) Examples of extracted XIC of $His_{10}$-SUMO1$^{T95R}$ modified peptides detected before and after K-ε-GG antibody enrichment. Bar plot represents the abundance of $His_{10}$-

SUMO1$^{T95R}$ modified peptides detected before and after K-ε-GG antibody enrichment.

(H) Examples of extracted XIC of His$_{10}$-SUMO1$^{T95K}$ modified peptide detected before and after K-ε-GG antibody enrichment. Bar plot represents the abundance of His$_{10}$-SUMO1$^{T95K}$ modified peptides detected before and after K-ε-GG antibody enrichment.

**Supplementary Figure 2. Comparison between SUMO1-conjugation sites identification in His$_{10}$-SUMO1$^{T95R}$ mutant cells, His$_{10}$-SUMO1$^{WT}$ mutant cells and His$_{10}$-SUMO1$^{\Delta GG}$ (related to Figure 1).**

(A) Bar plot shows the comparison among the total abundance of diGly modified peptides detected in His$_{10}$-SUMO1$^{T95R}$ cells, His$_{10}$-SUMO1$^{\Delta GG}$ and His$_{10}$-SUMO1$^{WT}$ cells.

(B) Examples of extracted XIC of diGly modified peptide detected in His$_{10}$-SUMO1$^{T95R}$ cells, His$_{10}$-SUMO1$^{\Delta GG}$ cells and His$_{10}$-SUMO1$^{WT}$ cells. The bar plot on the right shows the comparison of abundance of the diGly modified peptide detected in the three types of cell.

**Supplementary Figure 3. Analysis of SUMO1-conjugation sites and proteins in SUMO1 reference map and dynamic map. (related to Figure 2)**

(A) Venn plot shows the number of SUMO1-conjugation proteins detected in SUMO1 reference map, versus the number of SUMO1-conjugation proteins detected in SUMO1 dynamic map. Bar plots show the GO terms enriched by the proteins which were detected SUMOylated in SENPs overexpression.

(B) Venn plot represents the number of SUMO1-conjugation proteins detected in SUMO1 reference map, versus the number of SUMO1-conjugation proteins detected in each type of SENPs overexpressed cells. Bar plots show the GO terms enriched by the proteins which were detected SUMOylated in each type of SENPs overexpression.

(C) Bar plot shows comparison of the total number of SUMO1-conjugation sites detect in this study, and in Impens' study.

(D) Venn plot shows the number of overlapped SUMO1-conjugation sites/proteins detected in this study, and in Impens' study.

**Supplementary Figure 4. Representations of SUMOylated proteins and consensus motif in SUMO1 modified proteome (related to Figure 2).**

(A) Graphical representation of the location of SUMO1-modified sites for proteins MIB1, USP22 and NONO

**Supplementary Figure 5. DeSUMOylation patterns of SENPs mediated SUMO1 dynamic map (related to Figure 3) .**

(A) SubLogo representations of consensus motif in deSUMOylation sites. Asterisks marked the position of the SUMOylated lysine residue.

(B) Overview of GO enriched biological processes, cellular components and protein-domain families for potential SUMO1-deconjugated proteins.

**Supplementary Figure 6. The deSUMOylation function of SENP3 plays key role in RNA-virus induced signaling pathway (related to Figure 5) . .**

(A) Quantitative PCR measurement of *Ifnβ*, *ISG56*, *RANTES* and *IL-8* expression in HEK293 cells, HEK293 cells overexpression with the indicated plasmids after stimulation with 2μg/ml poly (I:C) 6hr. Graphs showed the mean ± s.d. and data shown were representative of three independent experiments. *$p < 0.05$; **$p < 0.01$ (two-tailed t-test)

(B) Loss of SENP3 attenuates poly (I:C)-induced anti-viral responsive genes. HEK293 cells transfected with the indicated plasmids and siRNA after stimulation with 2μg/ml poly (I:C) 6hr. Graphs showed the mean ± s.d. and data shown were representative of three independent experiments. *$p < 0.05$; **$p < 0.01$ (two-tailed t-test).

(C, D) SUMOylation of DHX15 and PCBP2 are elevated upon SENP3 depletion. HEK293T cells were transfected with the indicated siRNAs and plasmids. 40h post-transfection, cell lysates were subjected to immunoprecipitation and then immunoblotted with the indicated antibodies.

(E, F) Induction of *Ifnβ*, *ISG56*, *RANTES* and *IL-8* mRNAs were measured by quantitative PCR. HEK293 cells transfected with the indicated plasmids after stimulation with 2μg/ml poly (I:C) 6hr. Graphs showed the mean ± s.d. and data shown were representative of three independent experiments. *$p < 0.05$; **$p < 0.01$ (two-tailed t-test).

**Supplementary Table Legends**

**Supplementary Table 1. Matrix of SUMO1-conjugated proteome.**

Sheet 1. List of identified $His_{10}$-SUMO1$^{T95R}$-conjugated sites. Columns from left to right contain gene symbol, position of the SUMO-conjugated lysine residue in protein, amino acid sequence surrounding the SUMO site (-15; +15), SUMO1 detected in each condition.

Sheet 2. List of identified $His_{10}$-SUMO1$^{T95R}$-conjugated sites with PSM and abundance.

Sheet 3. List of identified $His_{10}$-SUMO1$^{T95R}$-conjugated sites detected by our study and published studies. Columns from left to right contain amino acid sequence surrounding the SUMO site (-15; +15), gene symbol, position of the SUMO-conjugated lysine residue in protein,

SUMO1 detected in each condition, SUMO1 detected in this study, and in other studies.

Sheet 4. List of identified $His_{10}$-SUMO1$^{T95K}$ -conjugated sites with abundance.

**Supplementary Table 2. GO enrichment analysis of SUMO1-conjugated proteome.**

Sheet 1. GO terms enriched by the SUMO1-conjugated proteome.

Sheet 2. GO terms enriched by the proteins that can be specifically deSUMOylated by SENP members.

**Supplementary Table 3. Matrixes of SENP specific deSUMOylated SUMO1-conjugated proteome.**

Sheet 1-6. Protein matrix of SUMO1-conjugated proteome specifically deSUMOylated by SENP members.

**Supplementary Table 4. GO enrichment analysis of proteins deSUMOylated by SENP members.**

Sheet 1. GO pathway analysis of proteins deSUMOylated by SENP members.

### Supplementary Figure 1

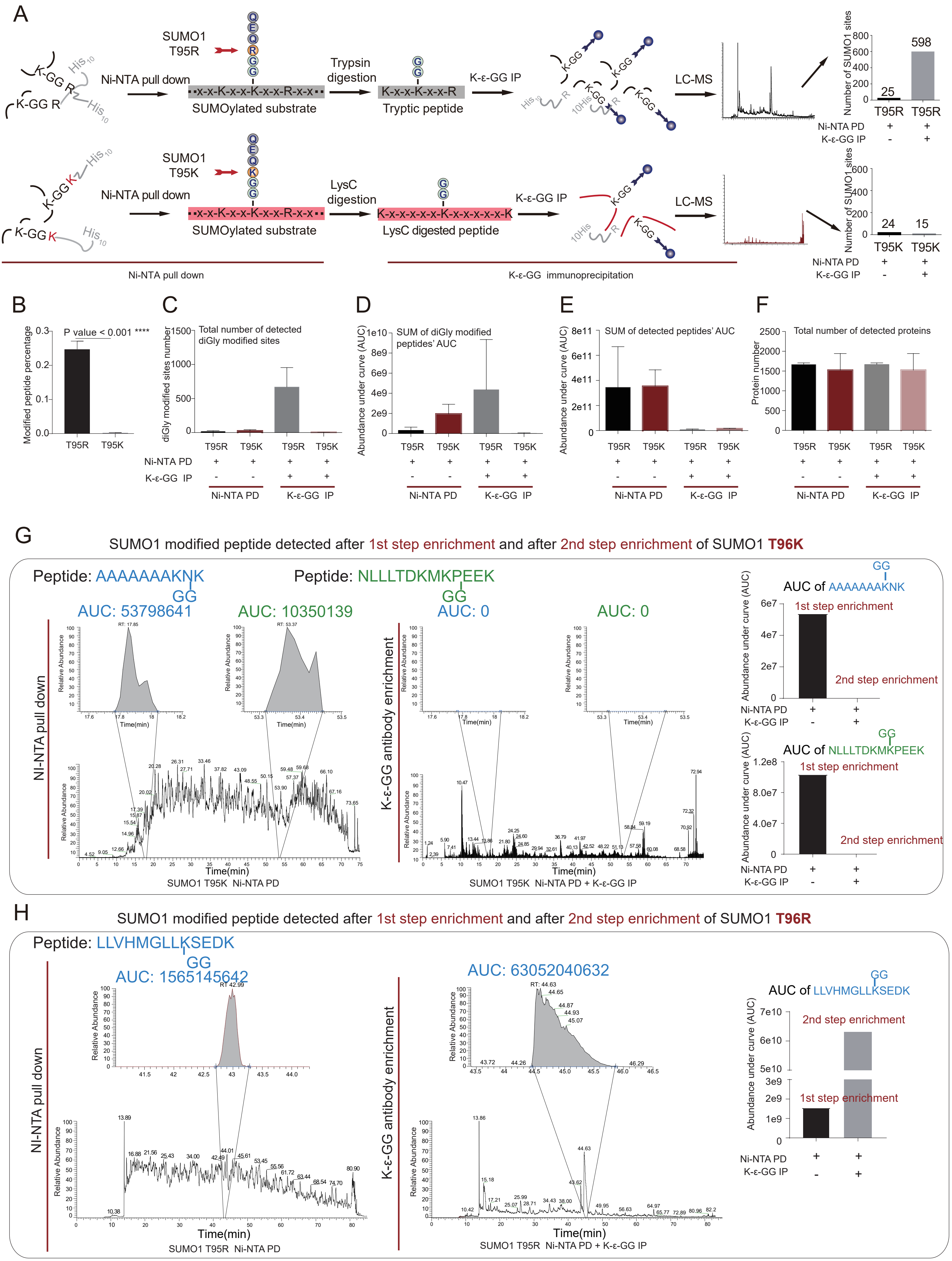

### Supplementary Figure 2

A

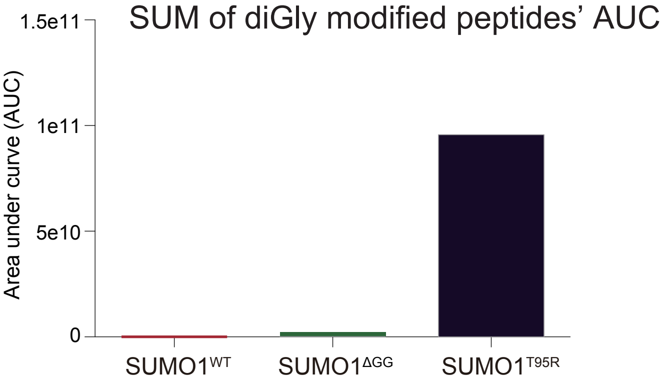


B

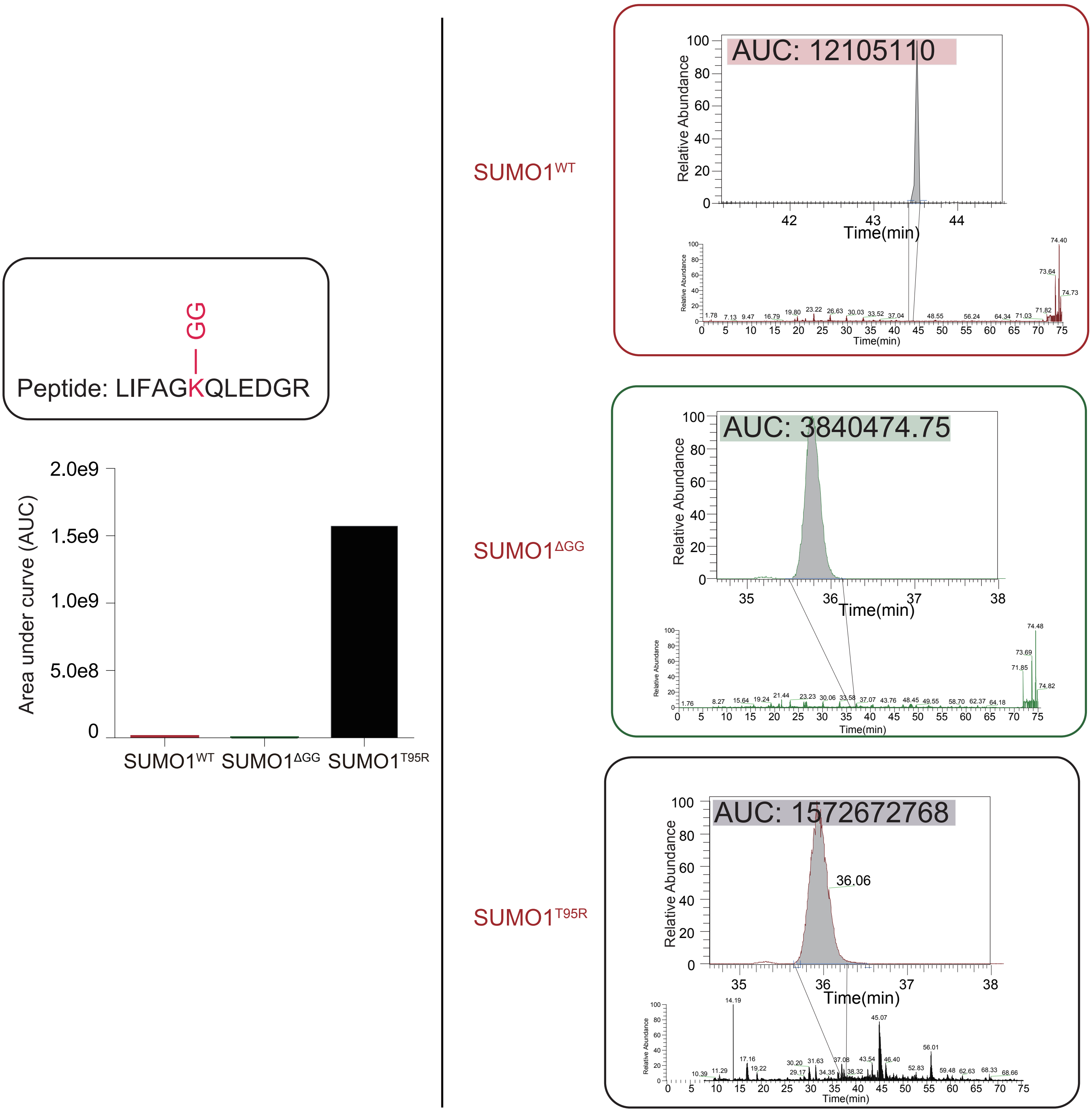

### Supplementary Figure 3

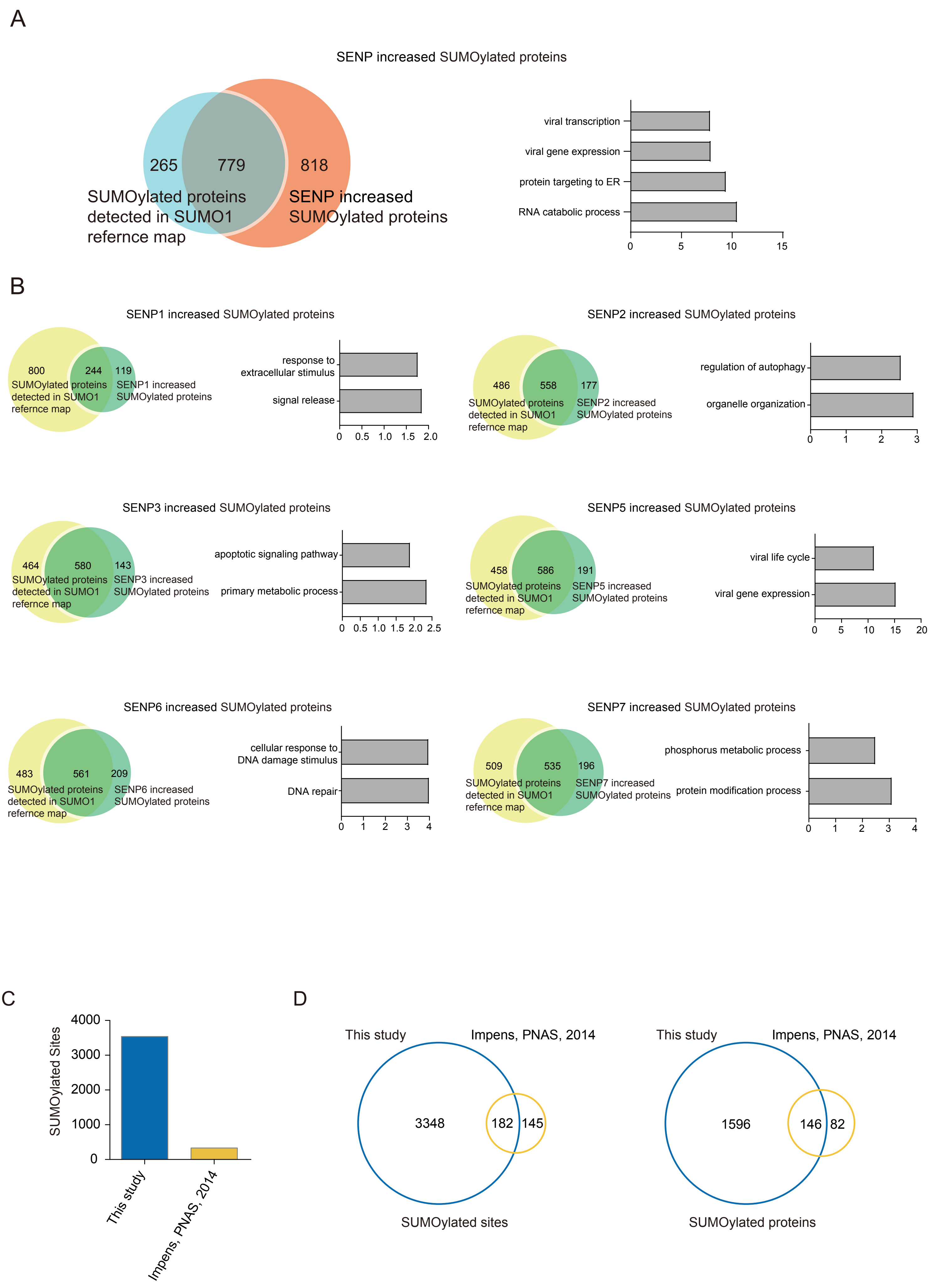

### Supplementary Figure 4

A

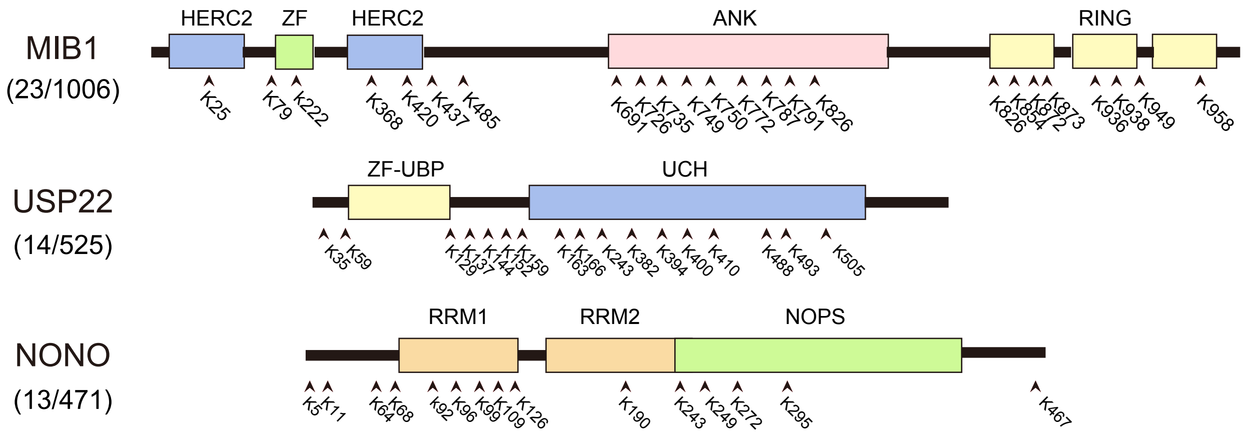

### Supplementary Figure 5

A

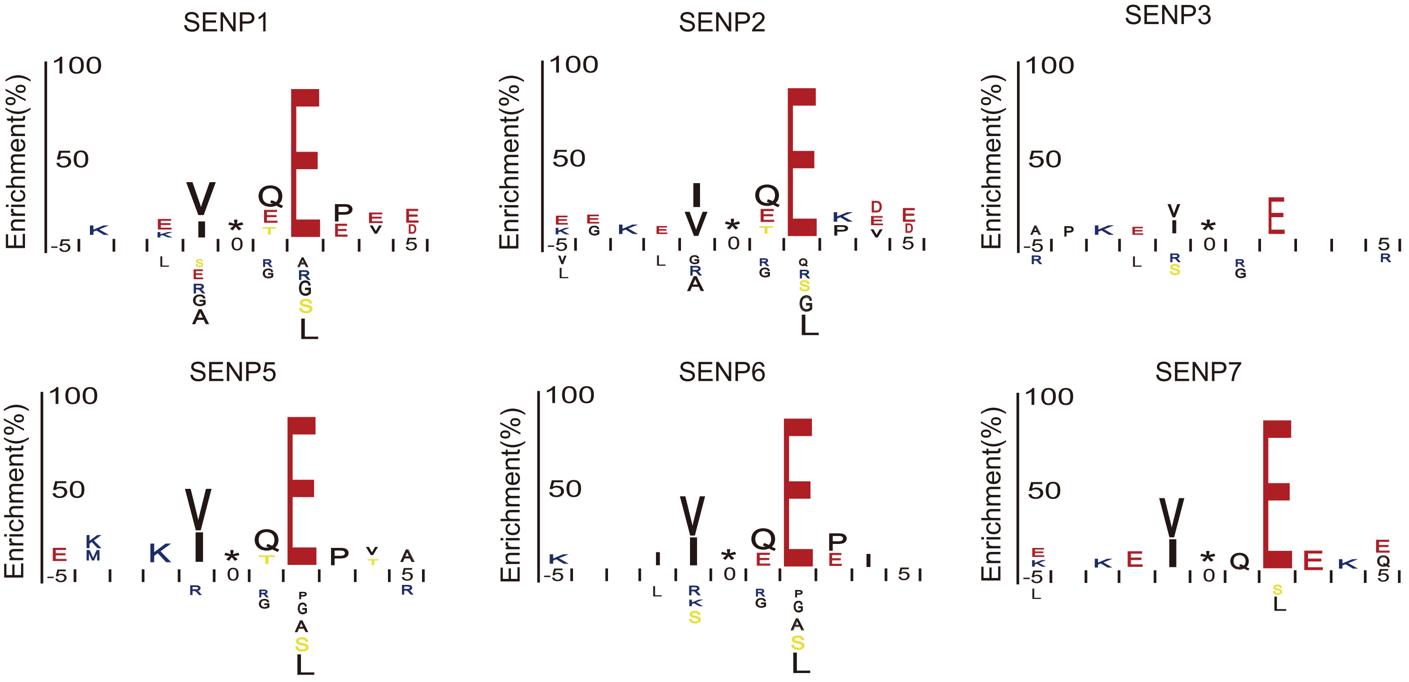


B

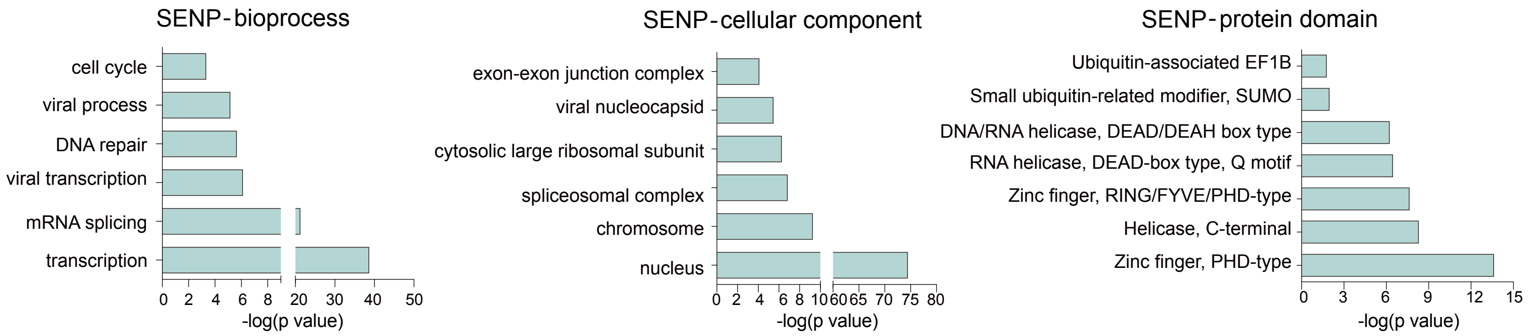

### Supplementary Figure 6

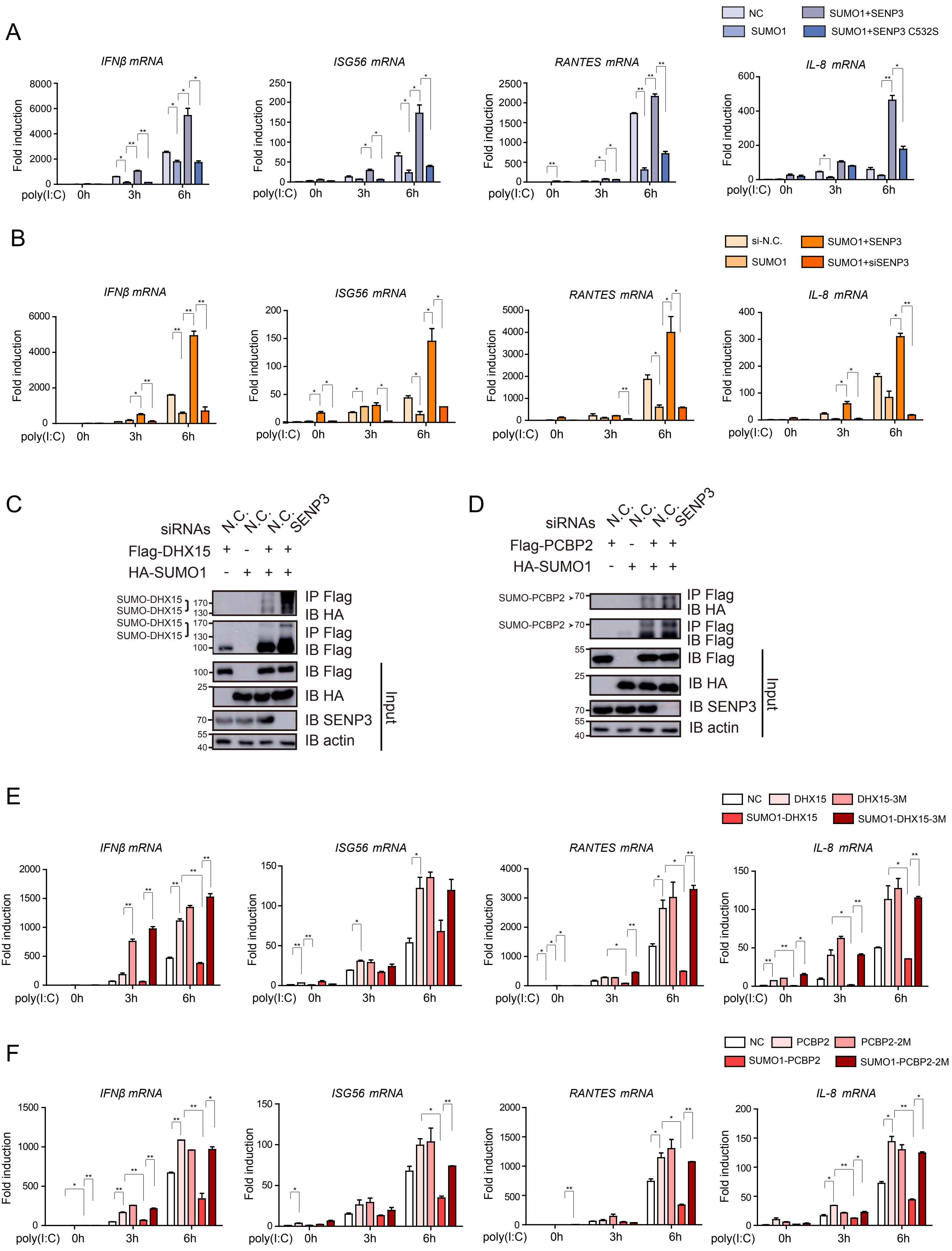
